# Motor context coordinates visually guided walking in *Drosophila*

**DOI:** 10.1101/572792

**Authors:** Tomás Cruz, Terufumi Fujiwara, Nélia Varela, Farhan Mohammad, Adam Claridge-Chang, M Eugenia Chiappe

**Author notes:** These authors contributed equally (Tomás Cruz, Terufumi Fujiwara).

## Abstract

Course control is critical for the acquisition of spatial information during exploration and navigation, and it is thought to rely on neural circuits that process locomotive-related multimodal signals. However, which circuits underlie this control, and how multimodal information contributes to the control system are questions poorly understood. We used Virtual Reality to examine the role of self-generated visual signals (visual feedback) on the control of exploratory walking in flies. Exploratory flies display two distinct motor contexts, characterized by low speed and fast rotations, or by high speed and slow rotations, respectively. Flies use visual feedback to control body rotations, but in a motor-context specific manner, primarily when walking at high speed. Different populations of visual motion-sensitive cells estimate body rotations via congruent, multimodal inputs, and drive compensatory rotations. However, their effective contribution to course control is dynamically tuned by a speed-related signal. Our data identifies visual networks with a multimodal circuit mechanism for adaptive course control and suggests models for how visual feedback is combined with internal signals to guide exploratory course control.

## Introduction

Locomotion is based on the coordinated action of multiple body parts that are typically controlled in a context-dependent manner (Bizzi et al., 1991; Foster and Higham, 2014; Sponberg et al., 2011). This context is highly variable, and originates from motor commands and other internal signals, as well as from the physical properties of the environment (Caggiano et al., 2018; Crapse and Sommer, 2008; Dickinson et al., 2000; Franklin and Wolpert, 2011; von Holst and Mittelstaedt, 1950; Sperry, 1950). Due to the variable nature of the context, locomotor performance is thought to depend on the interplay between self-generated sensory signals (sensory feedback), and motor-related internal information (Bässler and Büschges, 1998; Edwards and Prilutsky, 2017; Gordon et al., 2015; Tuthill and Azim, 2018). However, the general principles by which these multimodal and context-dependent interactions guide locomotion performance remain poorly understood.

Animals spend much of their time moving around extracting information about the environment. During exploration, flies and other animals display characteristic structured paths where frequent straight forward segments are interrupted by rapid changes in direction (Bell, 1990; Codling et al., 2008; Kim and Dickinson, 2017; Reynolds, 2018; Zeil, 1986). For visual animals, this particular locomotor strategy is advantageous to maximize acquisition of spatial information about the environment (van Breugel et al., 2014; Kim and Dickinson, 2017; Koenderink, 1986; Land, 1999; Muller and Wehner, 1988; Pfeffer and Wittlinger, 2016; Tolman et al., 1992). Maintaining a straight course, however, is a difficult task due to external or internal sources of noise (Dickinson et al., 2000; Franklin and Wolpert, 2011). This difficulty becomes more apparent when sensory feedback is perturbed, like attempting to walk on a straight line when blindfolded. Thus, an interesting possibility is that straightness performance depends on the interaction of visual feedback and other self-generated sensory and/or internal signals to generate an robust estimate of path deviations (body state), which can be used in the proper context for course control (Britten, 2008; Dickinson et al., 2000; Franklin and Wolpert, 2011; Körding and Wolpert, 2004; Pitkow and Angelaki, 2017; Todorov and Jordan, 2002).

The global structure of visual feedback consists of patterns of coherent retinal image flow, here simply described as “visual flow”, which results from movement of the body, head and eyes (Gibson, 1958; Koenderink, 1986). These complex patterns contain translational and rotational components that are related to the translational and rotational aspects of locomotion, respectively (Lappe et al., 1999). Rotational visual flow is thought to be important for controlling body rotations (Borst, 2014; Brandt et al., 1971; Srinivasan, 2011; Warren et al., 2001). Indeed, rotational visual flow is used by humans to detect small deviations from a straight course (Turano and Wang, 1994); however, its role in walking control remains largely contested (Cutting et al., 1992; Götz and Wenking, 1973; Harris and Bonas, 2002; von Holst and Mittelstaedt, 1950; Katsov and Clandinin, 2008; Prokop et al., 1997; Rushton et al., 1998; Warren and Hannon, 1988; Warren et al., 2001). This may be due to the additional contribution of other features of visual feedback to behavior (Koenderink, 1986). Cell-type specific recordings within networks processing visual flow in locomoting animals are critical for assigning the contribution of the different attributes and components of visual feedback to locomotor performance. Visual flow-sensitive neurons are found universally across species, from insects to non-human primates, and exhibit large receptive fields selective for specific patterns of visual flow (Britto et al., 1981; Duffy and Wurtz, 1991; Grasse and Cynader, 1982; Hausen, 1982a; Hengstenberg et al., 1982; Joesch et al., 2008; Kano et al., 1990; Krapp and Hengstenberg, 1996; Kubo et al., 2014; Morgan and Frost, 1981; Schnell et al., 2010; Soodak and Simpson, 1988). In non-human primates, visual flow-sensitive neurons combine visual and vestibular feedback to represent heading information (Bradley et al., 1996; Britten and van Wezel, 1998; Chen et al., 2011; Gu et al., 2008). However, neural recordings within this network have not yet been carried out in walking animals and therefore, it remains unclear what information might be represented during locomotion. To address this issue, and because of its small size, genetic tools, and existing knowledge of visual flow sensitive networks, we have performed such experiments in *Drosophila melanogaster*. This opened the opportunity to test the functional relation among rotational visual flow, the activity of rotational flow sensitive neurons and the control of rotations in walking flies.

Like primates, a class of fly rotational flow-sensitive cells, the HS cells, receive non-visual, rotation-related information (Chiappe et al., 2010; Fujiwara et al., 2017; Kim et al., 2015, 2017). During walking, visual and nonvisual information interact congruently to estimate the fly’s ongoing body rotations (Fujiwara et al., 2017). This result suggests that the fly brain uses visual feedback to control walking. In contrast, during flight, the nonvisual signals appear specifically in the context of body saccades (Schilstra and Hateren, 1999), and their interaction with visual signals is suppressive (Kim et al., 2017). Thus, HS cells, which project to premotor brain regions (Namiki et al., 2018), seem to be modulated in a locomotor-context dependent manner, a context defined either by walking vs. flight, or by a saccadic vs. a non-saccadic movements. The results from these studies pose an important question: what is the functional significance of the disparate multimodal modulation, specifically in relation to the control of visually guided locomotion? Here, we use Virtual Reality to examine whether the fly actively uses visual feedback to control locomotor performance in more naturalistic conditions, and which visual pathways may be involved in this control. In addition, we performed simultaneous recordings of neural and walking activity (Seelig et al., 2010) to identify the neural populations that could contribute to such control. By identifying leg-based saccades, we found two distinct motor contexts of the fly’s exploratory walking that are defined by the dynamics of the forward and angular velocities, and head-body movement coordination. We found a critical role of self-generated rotational flow for the control of the fly’s body rotations; interestingly, we found that this effect was conditional to the state of fly’s forward speed. We show that a concerted, multimodal activity within different populations of rotational flow-sensitive neurons represents body rotations. However, a forward speed related signal modulates their activity differentially. Some cells are enhanced while other cells are inhibited by walking speed, and this differential modulation matches the expected visual responses of the cells during forward walking. Remarkably, the contribution of rotational flow-sensitive neurons to body rotations is also tuned by the state of the fly’s forward velocity. Altogether, our data directly implicates activity in identified visual pathways to course control. Furthermore, our data suggests a general circuit mechanism that guarantees an adaptive contribution of different populations of neurons to course control, and suggests models for how sensory feedback is combined with internal signals in a motor-context specific manner to estimate the state of the body, and to control the performance of exploratory locomotion.

## RESULTS

### Walking speed and angular movements define distinct locomotor contexts

Exploratory flies walked in a 90mm circular arena with heated walls. A random-dot based visual stimuli was projected onto the arena’s floor, i.e., ventrally to the walking fly (**Fig. 1A**) (Katsov and Clandinin, 2008). We used real time tracking of the fly to update the position of the stimulus, such that it was “clamped” to the fly’s position (see **Methods**). Therefore, in this virtual world, the translational and rotational components of visual feedback were decoupled, and the fly generated visual feedback only by body rotations (**Fig. 1A,B**). Flies displayed typical exploratory walking paths, characterized by translations with high speed and fixed direction, and rapid rotations with spike-like angular velocity profile known as body saccades (**Fig. 1C,D, Movie 1**) (Geurten et al., 2014; Schilstra and Hateren, 1999; Zeil, 1986). Importantly, the lack of translational visual feedback does not appear to affect the structure of exploratory behavior.

**Figure 1.**
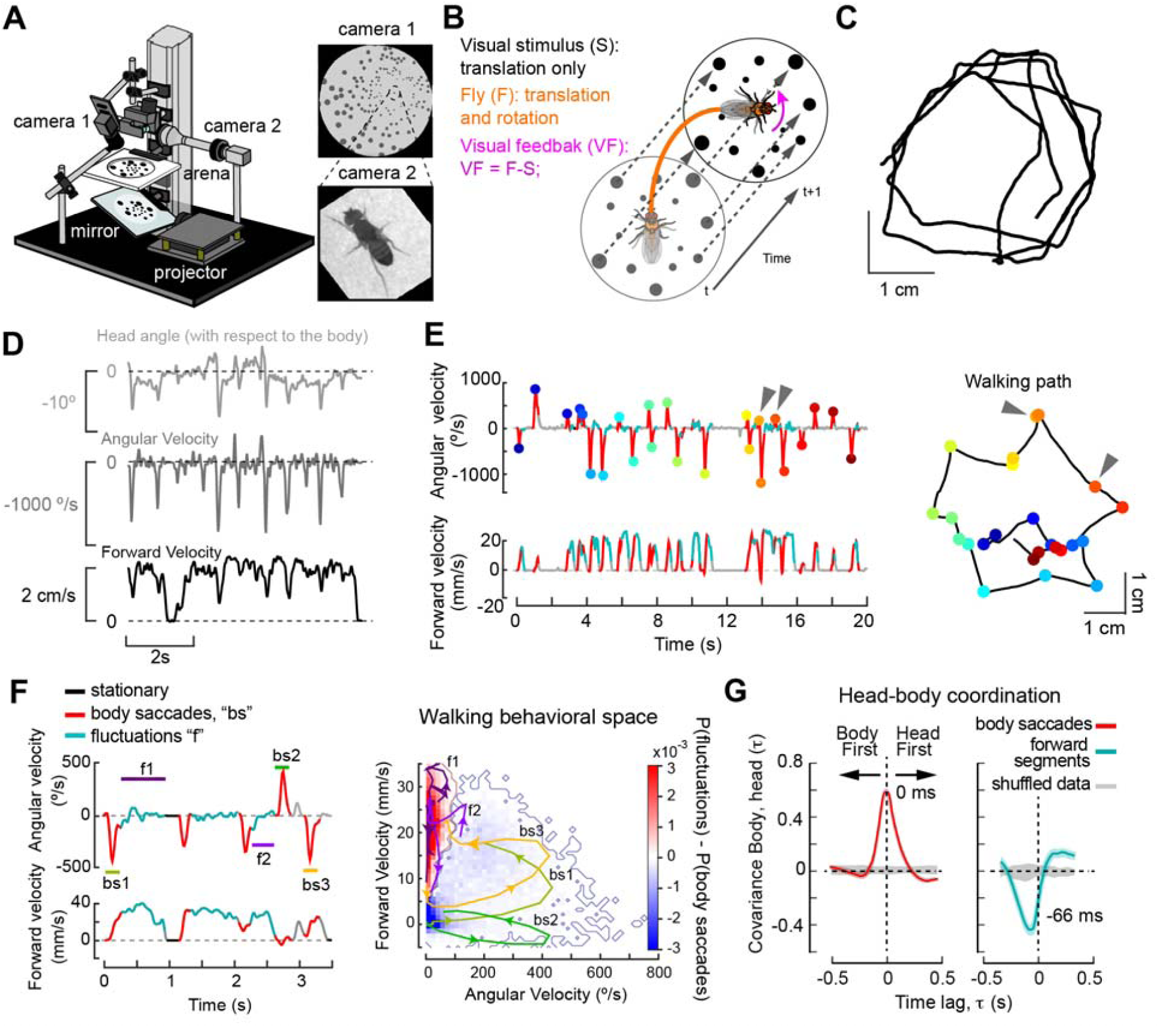
Exploratory walking in a virtual environment exhibits distinct motor contexts. (**A**) Virtual reality set up for freely walking flies. A pattern of random dots distributed radially from the fly was projected on to the floor of the arena. (**B**) A change in the fly position, detected in real time, elicited an identical translation of the stimulus, such that walking generated rotational visual flow only. (**C**) Example of exploratory paths of an individual fly. (**D**) Time series of angular velocity, forward velocity, and of head movements corresponding to the walking bout in (**C**). (**E**) Left, time series of a 20-s walking bout color-coded by identification of body saccades (red). Colored dots indicate individual saccades. Segments of high forward velocity without body saccades, denoted forward segments, are indicated in cyan (see **Methods**). Right, exploratory paths with body saccades highlighted in colors. Arrowheads highlight miniature saccades. (**F**) Left, Time series of a walking segment with body saccades (“bs”, red) and angular velocity fluctuations (“f, cyan). Right, Probability of angular velocity fluctuations over saccades as a function of the behavioral space of the fly. Color-code: red indicates regions with very low probability of body saccades, while blue indicates the reverse. Color-coded trajectories highlight different examples of trajectories of body saccades (bs1-3), and of separate forward segments (f1-2) within the behavioral space of the fly. (**G**) Head-body coordination measured as the cross-covariance between head angle and body angular velocity, during body saccades (left, red) or forward segments (right, cyan). Gray: cross-covariance analysis with temporally shuffled data.

The dynamics of body saccades were highly stereotyped within our dataset, and this property facilitated their identification through a dynamics-based classifier (**Fig. S1A, S2A,B; Fig. 1E**, see **Methods**) (Arthur et al., 2013). Our description of behavior was robust within the classifier’s parameter space (**Fig. S1B**). Body saccades were executed when the fly walked at low speeds (**Fig. 1F, Fig. S2A, E**), and the probability and amplitude of a saccade decreased with increasing walking speed (**Fig. S2E**). Notably, during these high-speed, non-saccadic walking segments, denoted “forward segments”, flies displayed slow angular velocity fluctuations with unstructured dynamics (**Fig. 1F, Fig. S2B**). Using an additional, high-magnification camera (**Materials** and **Methods**) (Bath et al., 2014), we observed that when flies executed a body saccade, the head and body rotated in the same direction with zero lag (at a frame rate of 120Hz), suggesting a shared motor program (**Fig. 1A, D, G**-left, **Movie 2**). In contrast, during forward segments, the head and body rotated in opposite directions, with the body preceding the head by a 66ms lag (**Fig. 1G**, right). These small-amplitude head movements vanished under head-fixation (**Fig. S2F**), suggesting that an artifact of the tracking system could not explain their presence. Interestingly, these small-amplitude head movements reduced gaze variability (**Fig. S2G, H**, p < 0.005 Wilcoxon signed-rank test, see **Methods**). Altogether, the analysis of body rotations and head-body movement coordination, suggests that body saccades and forward segments constitute two distinct locomotor contexts, which are associated with distinct states of forward and angular velocities, and different head-body movement coordination.

### Visual feedback controls forward segments but not body saccades

The prevailing view about the role of rotational visual feedback is that it informs the brain about body rotations to correct for deviations from an intended course. To study the relation between self-generated visual signals and walking performance, we analyzed the dynamics of body saccades and forward segments in darkness and compared them to saccades and forward segments in the presence of visual feedback. Body saccade dynamics were identical under visual feedback and darkness conditions, but their amplitude distribution was different (**Fig. S2B, C**). The lack of an apparent effect of visual feedback on saccade dynamics is consistent with previous work in tethered flight (Bender and Dickinson, 2006a; Heisenberg and Wolf, 1988). In contrast, forward segments were highly sensitive to visual feedback (**Fig. 2A**). To quantify this sensitivity, we calculated path straightness as a measure of course-control performance (**Fig. 2A**, see **Methods**). In darkness, flies walked less straight, and their path deviated more than in light conditions (**Fig. 2B**, left, p<10^-7^, MWW-Test, center), a result that is consistent with previous observations (Robie et al., 2010). That is, self-generated rotational visual signals seem to limit body rotations during forward segments. In addition, under visual feedback, flies also showed a tendency to increase slightly their speed, an effect that might be related to the control of path deviation (**Fig. 2B**, right).

**Figure 2.**
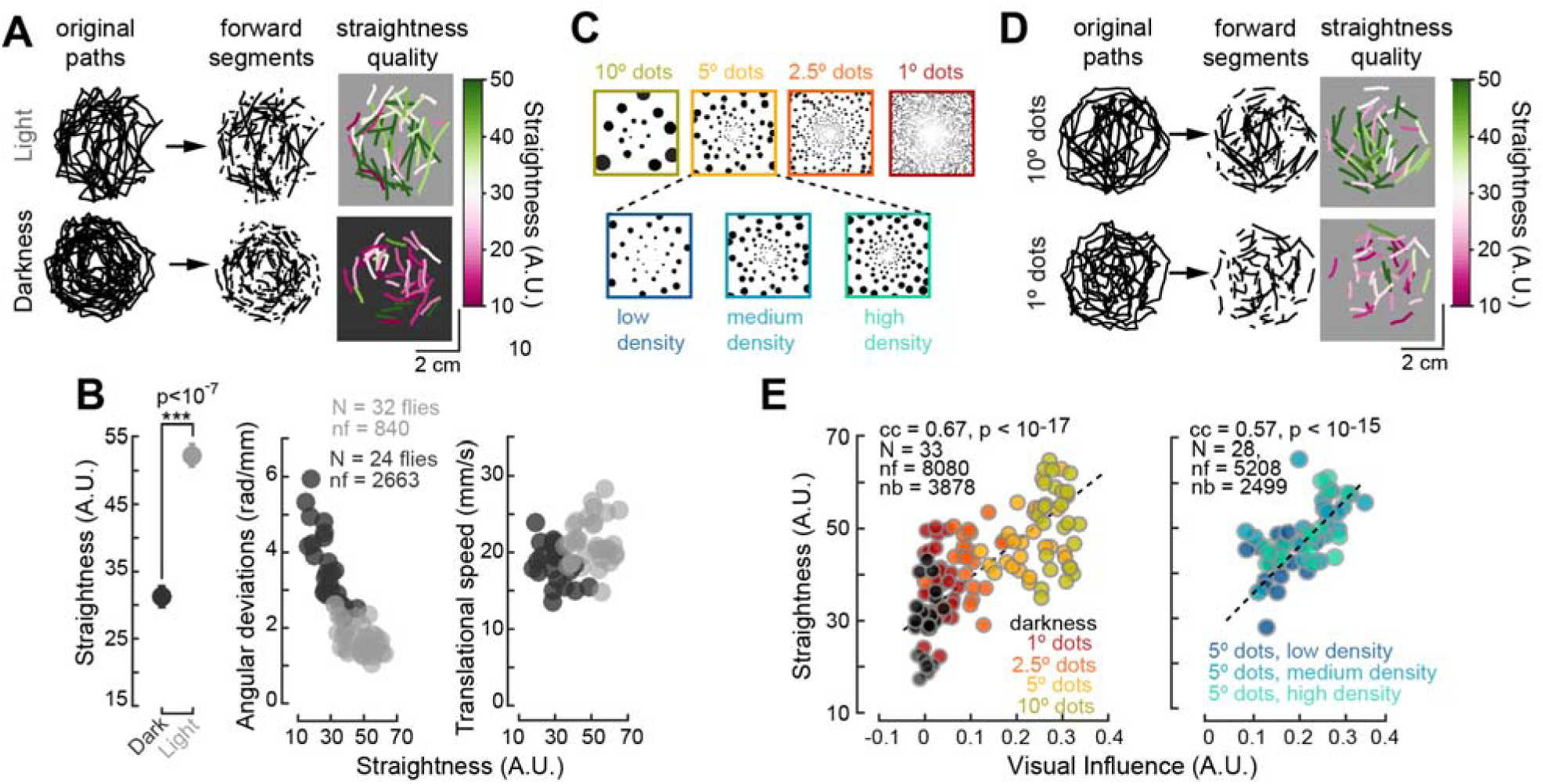
Visual feedback controls non-saccadic rotations during forward segments. (**A**) Exploratory paths of a fly walking with visual feedback (top) or in darkness (bottom). From the original paths (left), the classifier isolated fonvard segments (middle). The performance of these segments was measured by a straightness quality parameter (right, see Methods), with high values color-coded in greeri, and low values in magenta. (**B**) Left: mean straightness performance in darkness (dark gray) and light conditions (light gray) (±SEM, ***= p<10-7, MWW-Test). Middle: the cumulative angular deviation of the fly as a function of path straightness. Right: the translational speed of the fly as a function of walking straightness. (**C**) Stimuli with different physical aspects. (**D**) Similar to (**A**), example of a single fly’s exploratory paths under visual feed-back produced by different stimuli (in **C**) (**E**) Straightness performance as a function of visual influence, with size (left, p <10-17, t-test) or density (right, p< 10-15, t-test)as parameters of the physical aspect of the stimulus. For definition of visual influence, see main text, **Methods** and **Figure S3**.

To examine the relation between a fly’s sensitivity to self-generated visual stimuli and the individual’s walking performance, we designed a set of virtual environments with random dots of variable size or density (**Fig. 2C**). To evaluate the fly’s sensitivity to self-generated visual stimuli, we incorporated a “reversed gain” condition (see **Methods**). Under “natural gain”, a rotation of the fly in one direction makes the world rotate in the opposite direction, whereas under the “reversed gain” condition, the world moves in the same direction as the fly (**Fig. S3A**). If the fly detects the self-generated stimuli, reversed gain triggers a persistent rotation called circling behavior (von Holst and Mittelstaedt, 1950; Sperry, 1950) (**Movie 3**, **Fig. S3B, C**). The sensitivity of the fly to self-generated stimuli was defined as the difference in the probability of circling between the natural and reversed conditions, a scalar defined as “visual influence” (**Fig. S3D, E**, see **Methods**). A world with small dots (1° size) produced low visual influence, and flies in this world displayed straightness performance levels close to but better than those observed in darkness (**Fig. 2D, E**). On the other hand, flies exploring worlds with larger visual influence showed higher straightness (**Fig. 2D, E**). In fact, the more prominent the visual stimulus was, the more sensitive flies were to visual feedback, reflected in straighter walking paths (**Fig. 2E,** for dot size, p < 10^-17^; for dot density, p < 10^-15^, t-test). That is, the fly uses rotational visual feedback to minimize slow body rotations, thereby increasing the straightness of forward segments.

### Visual motion pathways underlie the control of body but not head rotations

What attributes of visual feedback underlie this active steering control during forward segments? Straightness control may rely not only on visual flow, but also on the local features of the visual feedback, a topic that has been a matter of debate (Cutting et al., 1992; Götz and Wenking, 1973; Harris and Bonas, 2002; von Holst and Mittelstaedt, 1950; Katsov and Clandinin, 2008; Prokop et al., 1997; Rushton et al., 1998; Warren and Hannon, 1988; Warren et al., 2001). Our data indicates that the global structure of the visual feedback contributed to the fly’s course control. First, the performance of the fly increased with increasing density of the dots, at least up to a certain limit (**Fig. 2E**, right). Such an effect of dot density on straightness performance would not be expected from a behavior based on a local stimulus features. Second, the averaged visual influence for stimuli with dot sizes below the acceptance angle of the fly eye (∼5°) was about two to four times stronger than in darkness (**Fig. S3E**) (Heisenberg and Wolf, 1984). Third, if a fly was monitoring a single dot (a local feature of the self-generated stimuli), we would expect the fly to follow it for some time, reflected on a bias in the individual’s orientation during exploration, but we found no evidence for such a bias (**Fig. S3F-I**). Fourth, the response amplitude of rotational flow-sensitive cells increased monotonically with the size of the dots (see below, and **Fig. S5**), suggesting that their contribution to the behavior may increase as a function of visual influence, which in turn correlates with straightness performance (**Fig. 2**). Altogether, our data indicates visual-flow processing pathways contribute to the control of non-saccadic rotations. In our experimental conditions, as in the real world, the local structure of visual feedback and associated visual pathways might additionally support straightness control.

Next, we asked whether motion vision was a critical cue for straightness performance. We used the Gal4-UAS system to selectively target the expression of the potassium inward-rectified Kir2.1 channel to silence the activity of T4/T5 cells (**Fig. 3A, B**), the first population of motion-sensitive and direction-selective cells in the fly visual system (Borst et al., 2010). Similar to control flies, experimental flies with silenced T4/T5 cells displayed characteristic exploratory paths (**Fig. 3C**). However, their forward segments were markedly less straight than those of control flies (**Fig. 3C, D** left). In fact, experimental flies lost their sensitivity to visual feedback (**Fig. 3D**, right). This insensitivity rendered course control performance close to darkness levels. Surprisingly, T4/T5 activity did not affect head-body coordination under natural gain (**Fig. 3E, F** “NG”), although the activity of T4/T5 cells did affect head-body movement coordination under visual perturbations, presumably due to the control of circling behavior, and/or to the head sensitivity to external visual motion stimuli (Kim et al., 2017) (**Fig. 3E,F**, “RG” and **Fig. S3I**). That is, self-generated visual motion is not critical for head-body coordination but fundamental for straightness performance, highlighting a differential role for rotational visual flow in gaze-vs. body-movement during exploratory walking.

**Figure 3.**
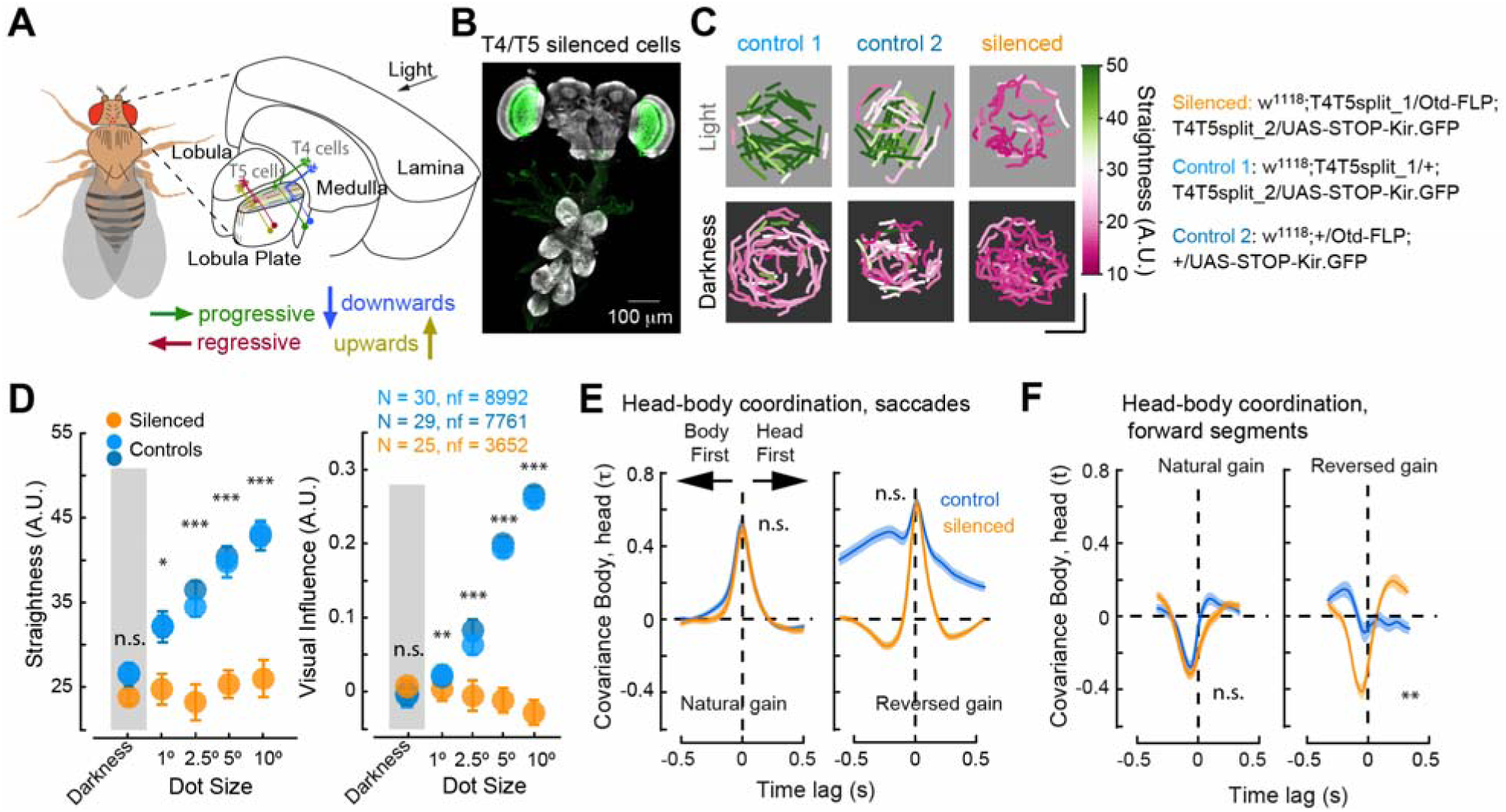
Visual motion is critical for straightness control but not for head-body coordination. (**A**) Schematic of the fly optic lobe highlighting the location of direction selective T4/T5 cells. (**B**) Expression pattern of Kir2.1::GFP targeted selectively to T4/T5 cells (green). (**C**) Exploratory paths of single flies with or without T4/T5 cells silenced (orange and blue colors, respectively. Visual environment: 10° dots. (**D**) Straightness performance (left), and visual visual influence (right) across different environments for control (blue) or experimental flies (T4/T5 cells silenced, orange). MWWTest, *p<0.05, **p<0.01, ***p<0.005 (**E**, **F**) Analysis of head-body coordination performed in control (blue) or experimental flies (orange), during saccades (**E**) or forward segments (**F**), and under natural (left), or reversed (right) gain conditions. **p<0.01, MWWTest.

### Contribution of progressive and regressive visual flow to the control of fly rotations

In the real world, a forward-walking fly generates a complex pattern of translational and rotational visual flow dominated by progressive (or front-to-back, FTB) visual motion (Lappe et al., 1999). However, in the virtual world with no translational visual feedback, a fly’s forward walking generates both FTB and regressive (back-to-front, BTF) motion that is induced by the slow rotations of the body (**Fig. 4A**). To test whether both directions contributed to straightness performance, we generated virtual environments where the stimulus was presented only to one eye (**Fig. 4Aii**). In this way, a stimulus projected onto the right eye in a fly rotating to the left would generate FTB visual flow only, whereas a fly rotating to the right will generate BTF visual flow only (note that the opposite is true under reversed gain). In these worlds, flies displayed the characteristic exploratory structure (**Fig. 1, 4B**), and the reversed gain condition induced circling behavior regardless of the direction of self-generated visual motion (**Fig. 4B, D**). However, the dynamics of the angular velocity during circling, i.e., under the RG condition, was different for FTB or BTF stimuli (**Fig. 4B,C**). FTB flow induced a large drift of angular velocity on top of which body saccades could be observed (**Fig. 4B**). This large drift coincided with moment when the fly walked at high forward speed (**Fig. 4C**), suggesting that it may correspond to the slow angular velocity fluctuations observed during forward segments under natural gain conditions. In contrast, BTF flow induced a rather small-amplitude drift, and more prominent body saccades in the same direction (**Fig. 4B**). That is, FTB and BTF pathways could jointly contribute to straightness performance; however, each pathway seems to control strength of angular fluctuations differentially in the context of stable, high walking speed (**Fig. 4C, D**).

**Figure 4.**
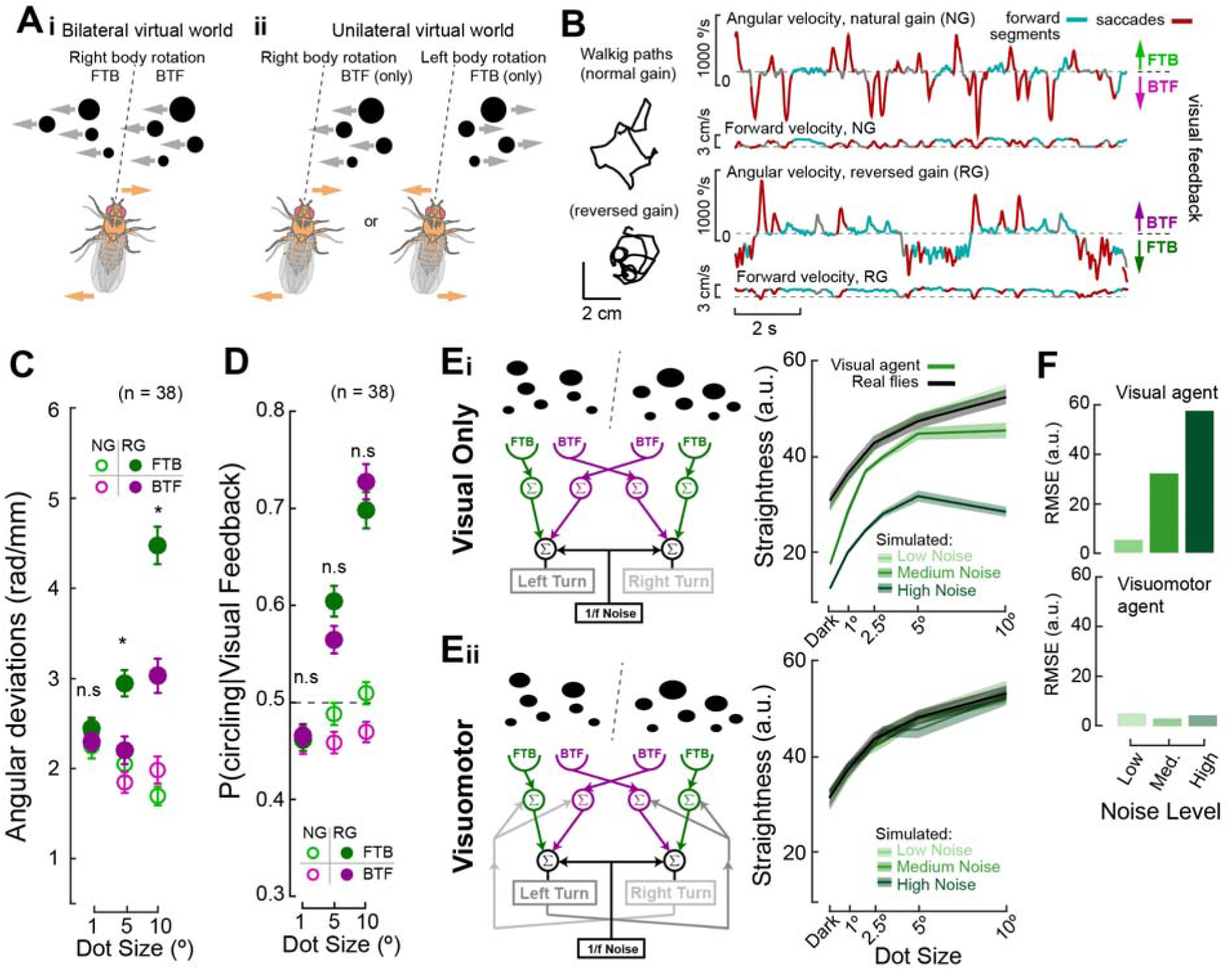
Progressive and regressive visual motion processing, and non-visual rotation-related information jointly contribute to robust course control. (**Ai**) In the VR world, a rightward rotation generates both Front-to-Back (FTB) and Back-to-Front (BTF) visual motion. (**Aii**) Under unilateral stimuli, a fly generates either direction of visual feedback. (**B**) Left, a walking-path segment of a fly in a unilateral 10°-dot size visual environment, under natural or reversed gain. Right, the angular (top trace) and forward (bottom trace) velocity time series of the corresponding example fly. (**C**) Mean angular deviations as a function of the visual environment’s dot-size. Grand-mean ±SEM, MWWTest, *p<0.05. (**D**) Conditional probability of circling given FTB or BTF visual feedback in different visual environments. Line represents chance. Grand-mean ±SEM, MWWTest. (**Ei**) Left, visual agent model schematic (see main text for explanation). Right, straightness performance of the agent (green color coded traces) and of the real flies (black trace) in different virtual environments, in the presence of different “noise” levels (hue code: light colors, “low noise’, dark color, “high noise”). (**Eii**) Same as in (**Ei**), but for a visuomotor agent. (**F**) Quantification of the residual difference between real fly and agent straightness performance for the different agent models in (E). RMSE: root mean squre error.

We built a model agent to test whether a simple control system based on FTB and BTF sensitive pathways would explain the fly’s behavior during forward segments (see **Methods**). Body saccades in this agent were triggered as a function of the distance to the wall (**Fig. S4A**). Visual flow sensitivity was modeled with four channels, each containing classical visual motion detectors (Borst et al., 2010). These four channels represented FTB and BTF systems, with the left FTB and the right BTF channels interacting to promote leftward rotations, and the right FTB and the left BTF channels interacting to promote rightward rotations (**Fig. 4Ei**). In addition, the model contained noise representing both internal and external sources (see **Methods**) (Dickinson et al., 2000; Franklin and Wolpert, 2011). Under the “low noise” condition, we tuned the two free parameters to approximate the agent’s behavior to the fly’s behavior (**Fig. 4E,** right**, F,** top; **Fig. S4B-E**). When the noise in the system increased, however, the visual agent’s course control performance degraded, and was markedly lower than the real fly’s straightness performance (**Fig. 4E,** right, **F**, top). FTB visual flow sensitive cells receive rotational-related non-visual signals (Fujiwara et al., 2017). We incorporated this finding to create a visuomotor agent (**Fig. 4Eii**, left). This visuomotor agent showed straight paths that matched the performance of real flies at all levels of noise (**Fig. 4Eii**, right, **4F**). A “motor-only” agent indicated that non-visual signals could explain course control of flies in darkness (**Fig. S4F**). Altogether, our data shows that a simple control agent could largely explain the behavior of the fly, and strongly suggests that multimodal interactions in such a simple control system increase course control robustness.

### Walking speed differentially modulates multimodal activity in flow-sensitive neurons

FTB- and BTF flow-sensitive neurons are located in separate layers of the fly Lobula Plate (LP) (Borst et al., 2010; Silies et al., 2014). HS cells, with dendrites in the first layer of the LP, display wide-field sensitivity to FTB visual flow (Barnhart et al., 2018; Chiappe et al., 2010; Hausen, 1982b; Krapp et al., 2001; Maisak et al., 2013; Schnell et al., 2010, 2012). In blowflies a class of wide-field, BTF-sensitive neurons with dendrites in the second layer of the LP—the H2 cell (Hausen, 1982b)—projects to a contralateral premotor area where it converges with axons of the HS cells (Borst, 2014; Hausen, 1984). We used this anatomical property to identify the *Drosophila* homologous H2 cell. Electroporation of Texas Red dextran at the axon terminal of HS cells revealed a single neuron projecting to the contralateral LP, with H2-like dendrites (**Fig. S5A,** see **Methods**) (Krapp et al., 2001). Whole-cell patch recordings from this spiking neuron showed BTF selectivity (**Fig. S5B**). Given these properties, we referred to this neuron as the likely H2 cell in *Drosophila*.

H2 and HS cells responded to moving random dots, with their response magnitude scaling as a function of the dot size (**Fig. S5B**). That is, H2 and HS cells were well posed to process self-generated FTB and BTF motion signals, and therefore, to contribute to the control of body rotations (**Fig. 3, 4**). Next, we tested whether H2 cells received rotational-related non-visual information, as is the case for HS cells (Fujiwara et al., 2017). In darkness, activity of H2 in walking flies confirmed the presence of such non-visual signals (**Fig. 5A,B**). The H2 cell was depolarized by ipsiversive fly rotations, whereas contraversive rotations inhibited it, and therefore, opposite side H2 cells have opposite non-visual direction selectivity (**Fig. 5A,B** and **Fig. S5C**). Similar to HS cells, this non-visual direction selectivity was angular velocity sensitive (**Fig. S5C**). To examine visual and non-visual signal interactions within H2 cells, we performed recordings of neural activity in a walking fly under visual stimulation. The visual stimulus was generated by the fly during a closed-loop trial but presented to the same individual in open-loop conditions (“replay trial”, see **Methods**). In this manner, we tested the neuron’s sensitivity to decoupled but simultaneously presented visual and non-visual signals (**Fig. 5C**). The direction selectivity of H2 and HS cells was highest when the neurons received congruent multimodal information, i.e. when by chance, visual stimuli was similar to the one that would have been generated by the fly’s own walking movements (**Fig. 5D,** and **Fig. S5D,** left, p < 0.05, two-way ANOVA followed by a Tukey-Kramer test). However, at high forward velocity, when multimodal activity of HS cells was boosted (**Fig. S5D**, right), H2 became insensitive to body rotations (**Fig. S5D**, right). The differential effect of forward speed on HS and H2 cells’ activity was evident under dark conditions. H2 was anti-correlated while HS was correlated with the fly’s speed, and these correlations were tightly coupled (**Fig. 5E**, HS cells: −6.5±19.5 ms, Mean±SD, n=18 cells; H2 cells: −2.9±68.5 ms, n=14 cells). Moreover, forward speed seemed to exert a multiplicative negative effect on H2’s non-visual selectivity: the faster the fly walked forward, the less responsive the cell became to the fly body rotations (**Fig. 5F**, left, slope difference between high and low speed p = 0.002, offset difference, p = 0.22, bootstrapping, see **Methods**). In contrast, forward velocity increased the overall activity of HS cells (**Fig. 5F**, right, slope difference, p = 0.792, offset difference, p = 0, bootstrapping). In summary, multiple, non-visual signals interacted with the visual responses of HS and H2 cells. Rotation-related multimodal information combines congruently in the two populations of cells, suggesting that these neurons could contribute in a concerted manner to a simple control system, as suggested by the visuomotor model agent (**Fig. 4**). However, this multimodal interaction is controlled by a speed-related signal in a cell-type specific manner; a manner that matches the expected response to FTB visual flow induced by forward walking, which excites HS but inhibits H2 cells.

**Figure 5.**
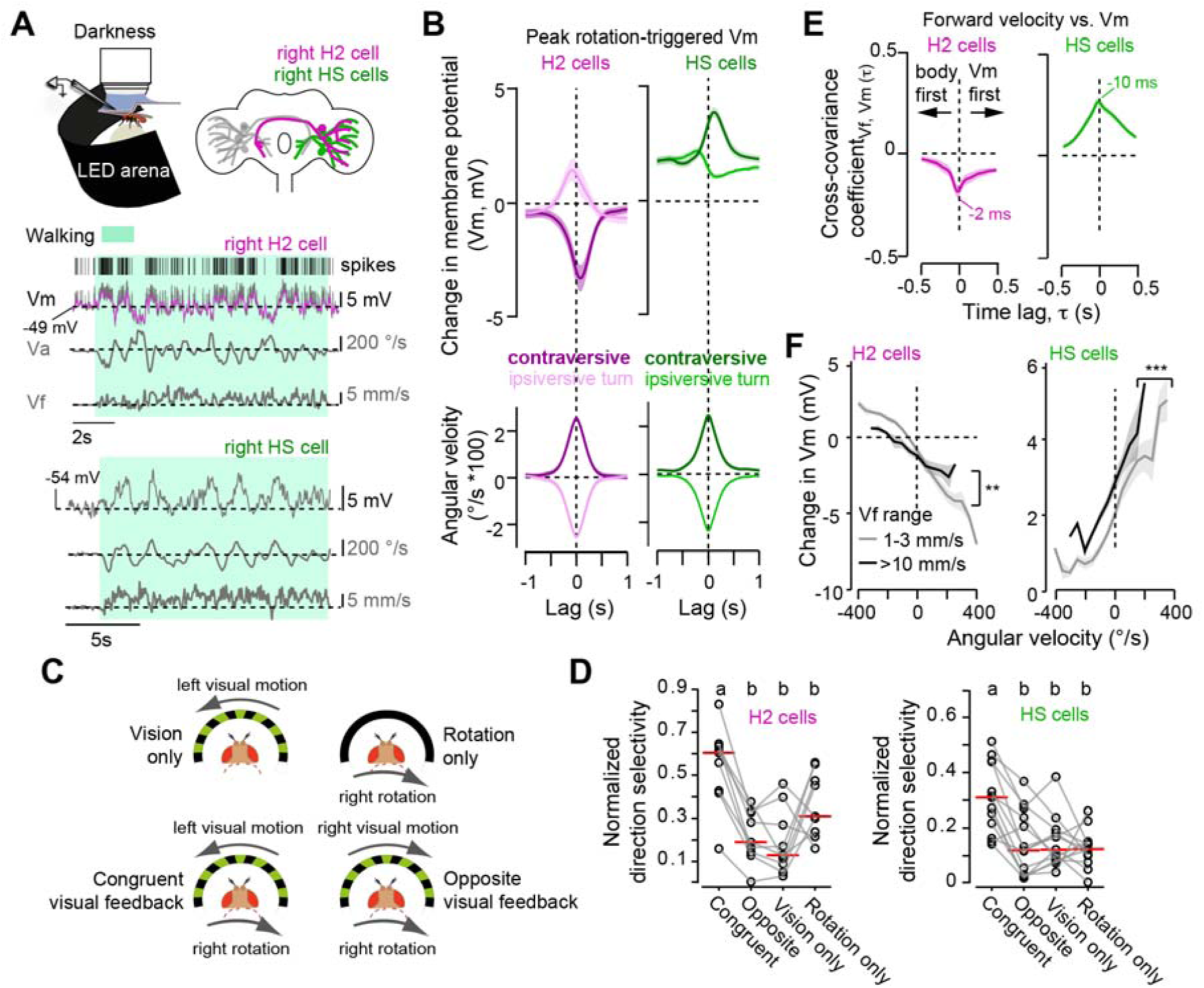
Congruent and motor context dependent visuomotor processing in HS and H2 cells during walking. **(A)** Top: schematic of the experimental setup (left), and of H2 and HS cells (right). Middle and bottom: Example segments of recordings in darkness of the cell’s membrane potential (Vm), and a fly’s angular (Va) and forward (Vf) velocities for a right-side H2 cell (middle), and for a right-side HS cell (bottom). The green shade indicates walking. For H2, the subthreshold Vm, and the spikes are indicated in magenta and with ticks, respectively. (**B**) Vm triggered at the peak of ipsiversive or contraversive fly rotations (bottom traces) for H2 (left, magenta, n=ie cells) and for HS cells (right, green, n=25 cells). (**C**) Four different visuomotor configurations during walking under visual stimulation. “Vision only” corresponds to visual stimulation during quiescence. “Rotation only” corresponds to moments when visual stimuli is stationary. “Congruent” is when a fly rotating to one direction received an opposite-direction visual motion stimulus. “Opposite” is when a fly rotating to one direction received same-direction visual motion stimulus. (**D**) Left, H2 direction selective responses to dynamic visual motion stimuli under different visuomotor configurations (mean ± SD, n=9 cells). Right, same for HS cells (right, mean ± SD n=13 cells). Red lines denote median values. Letters indicate significant difference among the different configurations (P<0.05, two-way ANOVA followed by Tukey-Kramer test). (**E**) Cross-covariance analysis between the fly’s forward velocity and the Vm of H2 (left, magenta. n=16 cells) or HS cells (right, green, n=25 cells) in recordings performed in darkness. (**F**) Change in Vm (relative to quiescence) as a function of the fly’s angular velocity during low (1-3 mm/s, light gray), or high (>10 mm/s, dark gray) forward speed for H2 (left, n= 9 cells) or HS cells (right, n= 9 cells). **: p = 0 002 for slopes, and ***: p= 0 for offset, boostrapping method (see **Methods**). In (B) and (E, F), data is presented as the mean ± SEM with each cell recorded from a separate fly.

Critical to the control system is the direction selectivity of FTB- and BTF-sensitive neurons (**Fig. S5B, 4**). When a fly rotates, this direction selectivity promotes unbalanced activity within the bilateral neural population. Unbalanced HS activity across the brain hemispheres induces a bias in the direction of walking flies (Busch et al., 2018; Fujiwara et al., 2017). Because H2 cells converge with HS cells’ axons, we predicted that unbalanced activity within the bilateral population of H2 cells would also promote a bias. We selectively expressed the vertebrate ATP-gated cation channel, P2X_2_, in HS and H2 cells (**Fig. S6A-C**, see **Methods**). Confirming previous results (Fujiwara et al., 2017), we observed that unilateral activation at the axon terminals of HS cells promoted an ipsilateral bias in the angular velocity of the fly (**Fig. 6A**, p < 10^-3^, Wilcoxon signed-rank test). Interestingly, this manipulation also triggered an increase in forward speed (p = 0.01, Wilcoxon signed-rank test). Similarly, unilateral activation of H2 at its axon terminal induced a bias in fly rotations towards the injection site (**Fig. 6A**, p < 10^-3^, Wilcoxon signed-rank test) with an increase in forward velocity (p < 0.01, Wilcoxon signed-rank test). Unilateral activation at the dendrites of H2 cells, which are located within the opposite side hemisphere, induced a bias in the fly’s angular velocity contralateral to the site of stimulation (p < 0.01, Wilcoxon signed-rank test) with a marginal increase in forward speed (p = 0.037, Wilcoxon signed-rank test). Next, to test whether the observed effect was related to the convergent projection of HS and H2 cells, we performed a similar experiment in VS cells, a population of visual flow-sensitive neurons with projections in a different region of the premotor area (Borst, 2014; Hausen, 1982b, 1984; Namiki et al., 2018). Unilateral activation at the axon terminals of VS cells (**Fig. S6C**) did not induce a significant bias in the angular velocity of the fly (**Fig. 6A**, p = 0.08, Wilcoxon signed-rank test). That is, the effect of HS and H2 cell may be specific to premotor networks that receive concerted information from these two populations of cells. These results suggest that HS and H2 cells could together contribute to the control of the fly’s slow body rotations during forward segments. However, given that fly speed modulated HS and H2 cell activity differentially, the effective contribution of either population may vary dynamically and conditionally given a specific motor-context.

**Figure 6.**
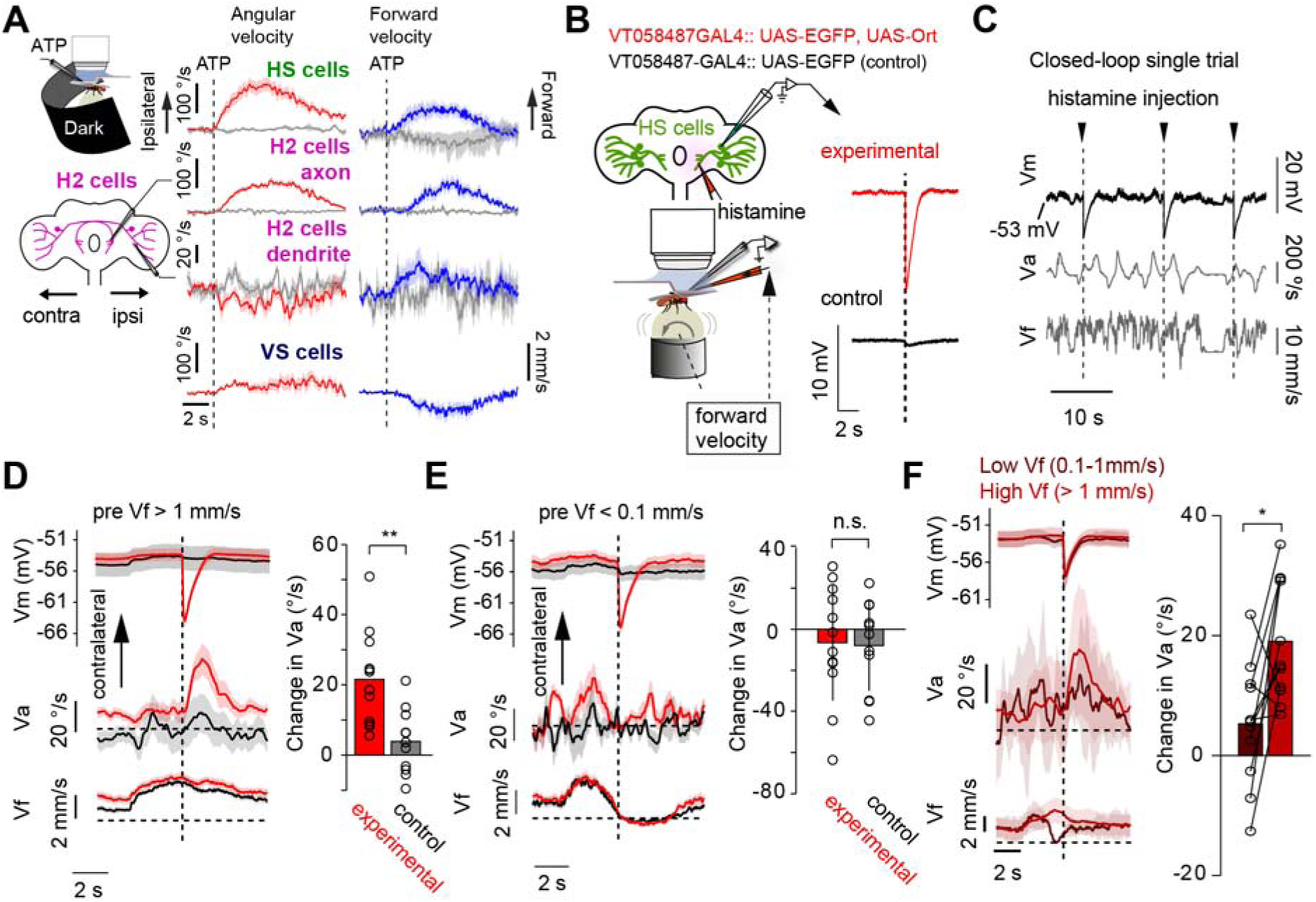
Motor context-dependent contribution of HS cells to course control. **(A)** Unilateral application of ATP and its effect on the angular (Va) and forward (Vf) velocities of a walking fly. Top: ATP application at the axon terminate of JIG cells (N-18 flies); middle, either at the axon terminals (N-14 flies), or at the dendrites (N=10 flies) of H2 cells; bottom: at the axon terminals of VS cells (N=10 flies) (mean ± SEM, 10 averaged trials per fly). Black traces indicate ATP application in control flies (N=10 flies) that did not express P2X2. **(B**) (Left) Experimental paradigm. (Right) The membrane potential (Vm) of HS cells (mean ± SEM, 10 averaged trials per fly) upon local application of histamine in experimental (red, top, N = 12 flies) or control flies not expressing ort artificially (black, bottom, N= 10 flies). (**C**) Example segment of simultaneously recorded Vm, Va and Vf. The arrowheads and dashed lines indicate conditional histamine injections. (**D**) (Left) The mean ± SEM of the Vm (N= 7 for experimental flies, N = 6 for control flies), Va and Vf in experimental (red, N=12 flies) or control flies (black, N=10 flies) upon conditional histamine injections. Note that Vm is reported tor a subset ottlies in which the whole-cell condition lasted until the end of the experiment. (Right) Change in Va before and after histamine injection in experimental (red) or control flies (black). **: p = 0.002, Wilcoxon rank-sum test. (**E**) Similar to (**D**) with the analysis performed in trials with average Vf less than 0.1 mm/s. (**F**) Similar to (**D**) with the analysis divided into low (dark red) or high (light red) Vf for experimental flies (N=11 flies). *: p = 0.02, Wilcoxon signed-rank test.

### Walking speed dynamically controls the contribution of HS cells to body rotations

It then follows that HS cells would have the strongest contribution to behavior during high forward speed, when H2 contribution would be minimal (**Fig. 5E, F, S5E**). We tested this idea by silencing the activity of HS cells with the targeted expression of Kir2.1 in HS (and in three additional VS cells (VS3-6) that were also labeled) using an intersectional strategy (**Fig. S6D, S7A**). Chronic bilateral inhibition of the activity of HS cells (mean resting potential, Vr, in experimental flies = −80.0±4.2 mV, mean ± SD, n=4 cells; Vr in control flies = −52.8±6.2 mV, n=5 cells) affected neither straightness performance, nor head-body coordination (**Fig. S7**). This result suggests that other parallel networks might contribute to straightness performance (Hausen and Wehrhahn, 1990), that HS activity might not contribute to the control of the slow rotational fluctuations, or that the control system might respond to unbalanced activity within the bilateral populations of neurons (**Fig. 6A**), as suggested by our model (**Fig. 4**). To test this last idea, we used opto- and chemogenetics to reversibly and temporally silence the activity of HS cells unilaterally and conditionally based on the state of forward speed. We expressed the light-gated anion channel GtACR1 in HS cells (**Fig. S6D**) (Busch et al., 2018; Govorunova et al., 2015; Mohammad et al., 2017), and activated the channel once the fly increased the forward velocity passing a threshold (**Fig. S6E-G,** see **Methods**). HS cells expressing GtACR1 showed robust, intensity-dependent hyperpolarization upon light illumination, whereas HS cells without GtACR1 showed a slight depolarization, presumably due to visual, or thermal effects related to the linear absorption of light (Stujenske et al., 2015) (**Fig. S6E**). Under these experimental conditions, the conditional unilateral inhibition of HS cells caused an initial rapid contralateral rotation to the onset of light, followed by a sustained contralateral bias in angular velocity (**Fig. S6F**). Upon termination of the stimulation, flies reacted with an ipsilateral turn. Control flies displayed the same onset and offset responses, which indicated the presence of an artifact induced by the light illumination. However, their behavior showed a smaller and more variable bias in angular velocity upon stimulation (**Fig. S6G**). In a set of parallel experiments, we expressed the native histamine-gated chloride channel *ort* in HS cells, and locally applied histamine at the axon terminals to inhibit the neurons once the fly increased the forward velocity passing a threshold (see **Methods, Fig. 6B, C**) (Liu and Wilson, 2013). Control flies with no artificial expression of *ort* showed minimal inhibition upon histamine application, indicating that HS cells do not express high-levels of *ort* endogenously. In contrast, a brief pulse of histamine (10 ms, 1 mM) induced a reliable hyperpolarization in HS cells expressing *ort* (−11.9±0.72 mV at the peak, mean ± SEM., N=12 cells, **Fig. 6B, C**), and this perturbation caused a contralateral bias in the angular velocity of experimental, but not control, flies (**Fig. 6D** p < 0.01MWW test). Notably, despite the fact that HS cells were inhibited consistently and robustly, the amplitude of the effect on behavior depended on the state of the forward speed of the fly; at very low speed, the effect was negligible (**Fig. 6E**, p = 0.88, Wilcoxon signed-rank test), but increased as walking speed increased (**Fig. 6F**, p = 0.02, Wilcoxon signed-rank test). That is, unbalanced activity within the bilateral population of HS cells induced a slow compensatory bias in the angular velocity of the fly, especially prominent when the state of forward speed was high. This high walking speed on the ball is likely related to forward segments during freely exploratory walking, with the observed rotational biases likely representing a compensatory movement in response to a detected angular deviation of the fly.

## Discussion

The external presentation of rotational visual stimuli elicits robust eye, head or body rotations that can be induced even if individuals are stationary. However, the role of the underlying neural circuits driving these so-called compensatory movements in more naturalistic contexts, such as in the presence of self-generated visual signals during locomotion, have remained largely elusive. Here, we first describe two distinct, internally driven motor contexts for exploratory walking that are defined by the state of the forward speed, the dynamics of the angular velocity of the body, and the concomitant head movements (**Fig. 1,2 S1-2**). Second, we show that the fly actively uses visual feedback to control walking performance. Self-generated visual motion cues are critical for the control of the body’s rotations, but not for the head’s yaw rotations, and, interestingly, this control is implemented in a motor context-dependent manner (**Fig. 2,3, S2**). Our data further indicates that a concerted and multimodal activity of a network of rotational flow sensitive cells robustly monitors body rotations (**Fig. 4-5**). Strikingly, we found that the state of the forward speed dynamically gates the contribution of different elements within the network to course control by boosting the activity in some cells, while inhibiting others (**Fig. 5, 6**). Altogether, these results reveal a circuit mechanism that is based on the interaction of multimodal signals to adaptively control the straightness performance of an exploratory fly (**Fig. 7**).

**Figure 7.**
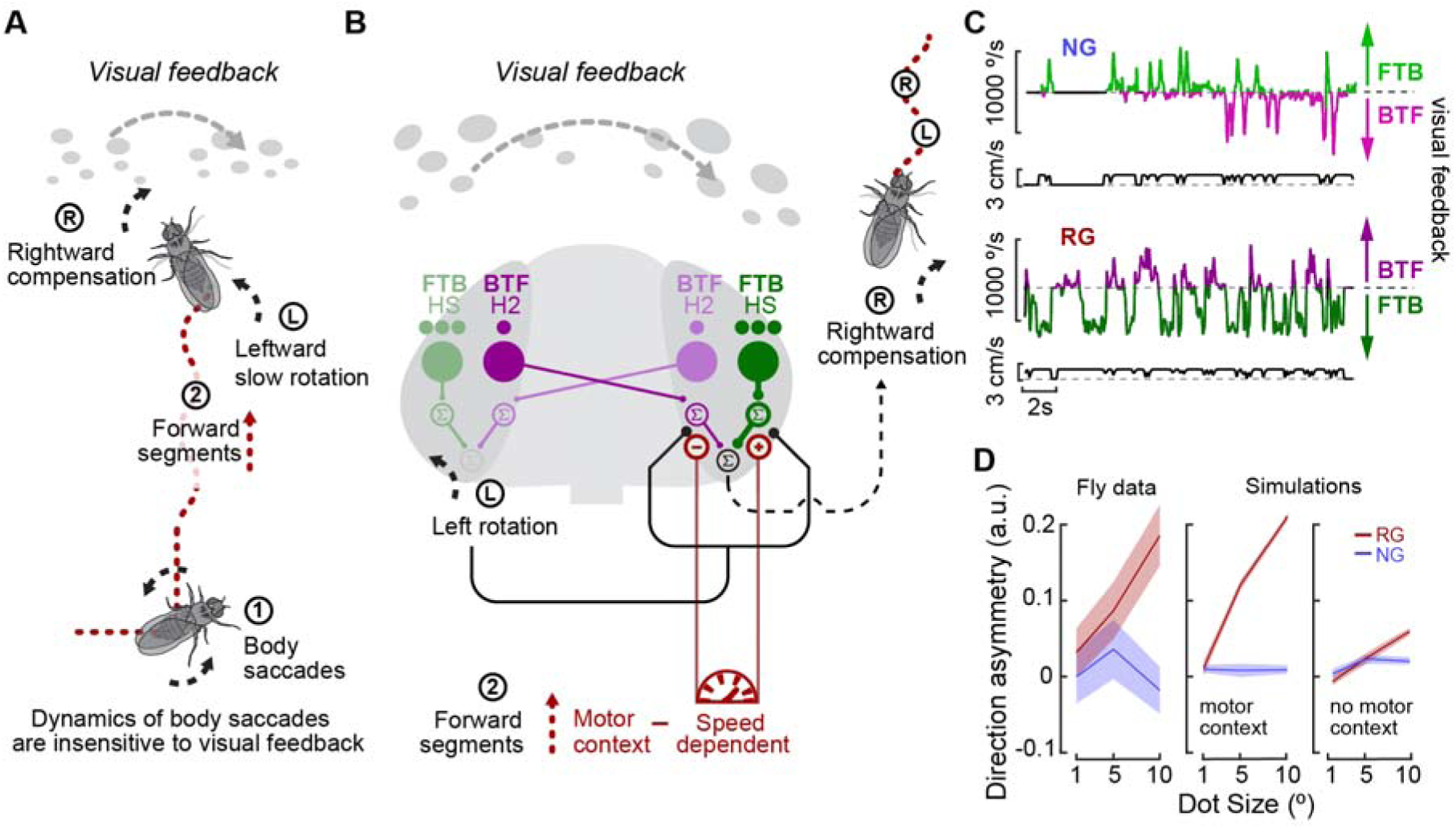
Motor context dynamically coordinates the concerted multimodal activity of visual flow-sensitive cells for course control. **(A)** Exploratory walking. Schematic highlighting the two distinct motor contexts, body saccades (1}, and forward segments (2) of the fly’s exploratory behavior. (**B**) Dynamic, multimodal course control system for straightness performance A rotation to the left during forward walking (L) induces visual feedback and non-visual rotation-related signals. This multimodal information is combined in FTB and BTF channels, represented by HS and H2 cells, to promote a compensatory rotation to the right. Dots below HS and H2 cells represent state of activity (1 =low, 3=high) induced by visual feedback during forward walking. An internal speed related signal, which is in a high state during forward walking, dynamically tunes the concerted contribution of BTF and FTB pathways to compensatory rotations. (**C**) Visuomotor agent simulation with an additional speed-dependent gating of the FTB and BTF channels under unilateral visual stimulus (and with an intermediate noise level, see Figure 4Eii). Tup. natural gain, “NG”, Bottom, reversed gain, “RG”. (**D**) Direction asymmetry calculated by the difference in mean angular fluctuations under BTF and FTB for the behavior data (Left), a model that includes motor context modulation (middle) and a model without it (right).

### Task-specific structure of exploration and context-dependent visual feedback control of walking

In this study, we examined the role of visual feedback in the control of saccades, and in the capacity of the fly to move in a fixed, straight course. We took an approach guided by the saccade-fixation structure of exploratory locomotion, and by task performance (Harris and Wolpert, 1998; Pitkow and Angelaki, 2017; Todorov, 2004) rather than by aiming at a full description of the modes of walking an exploratory fly may take (Berman et al., 2014; Kabra et al., 2013; Katsov et al., 2017). Although flies can initiate miniscule body saccades while moving forward (**Fig. 1E**, arrowheads, **S2E**), typically these rapid changes in walking direction are accompanied by stops in translation (**Fig. 1, Fig. S2**) (Blaj and Van Hateren, 2004; Geurten et al., 2014). The most prominent saccades are triggered close to the hot walls of the arena and are associated with directed backwards walking (**Fig. S2A, D**), likely corresponding to evasive maneuvers. As in flight (Censi et al., 2013; Muijres et al., 2015), the dynamics of these evasive maneuvers appear identical to other lower amplitude saccades (**Fig. S1A,B**). Regardless of their vigor, and as previously reported for tethered flight (Bender and Dickinson, 2006a; Heisenberg and Wolf, 1988), once the command for saccade is triggered, visual feedback does not contribute to the control of this rapid maneuver (**Fig. S2B**). Two potential mechanisms may explain the lack of visual sensitivity. First, the velocity-tuning curves of visual neurons may not be sensitive to visual stimulus velocity self-generated by a saccade. Second, recent studies in tethered flying flies have shown that populations of visual flow sensitive neurons receive precise, non-visual inputs that cancel responses to saccade-induced visual signals (Kim et al., 2015). Given that HS cells can respond to external visual motion stimuli with saccade-like velocity (Kim et al., 2017), we propose that leg-based saccade insensitivity to visual feedback may stem from similar neural suppressive mechanism.

The dynamics of the angular velocity fluctuations observed during forward segments is unstructured and largely composed of a unilateral drift (**Fig. S2B**), suggesting that these fluctuations do not simply correspond to the periodic walking stride cycle (Kress and Egelhaaf, 2012, 2014; Mendes et al., 2013; Strauss and Heisenberg, 1990; Wosnitza et al., 2013). We found that self-generated rotational flow controls the amplitude of the fluctuations (**Fig. 2,3**). Without this control, flies deviate from a straight course (**Fig. 1F, 2**), directly demonstrating the contribution of rotational flow to course control, an idea that has been contested (Cutting et al., 1992; Götz and Wenking, 1973; Harris and Bonas, 2002; von Holst and Mittelstaedt, 1950; Katsov and Clandinin, 2008; Prokop et al., 1997; Rushton et al., 1998; Warren and Hannon, 1988; Warren et al., 2001). Altogether, our data shows that moving forward and turning rapidly are based on separate, likely mutually exclusive, motor programs (**Fig. S2A, E**), which define distinct locomotive internal motor contexts (**Fig. 7A**). During forward segments, the motor commands arriving to the ventral nerve cord are based on internal signals driving forward speed, as well as on compensatory signals that maintain straightness and that depend on visual and other sensory feedback.

### Visual feedback is not critical for gaze stabilization

Saccades curtail the time spent generating retinal image shifts as a fly changes the direction of locomotion. We confirmed previous observations, both in flight and in walking, that the head and the body of the fly move in unison during a saccade (**Fig. 1, Fig. S2**) (Blaj and Van Hateren, 2004; Geurten et al., 2014; Heisenberg and Wolf, 1979; Schilstra and Hateren, 1999; Zeil, 1986). Like the rapid body maneuvers, head movements during saccades are insensitive to visual feedback (**Fig. 3**). Similarly, and despite the fact that head rotations are sensitive to external visual perturbations (**Fig. 3E “RG”, F “RG”, S3I**) (Kim et al., 2017), when flies walk forward, head-body movement coordination is largely unaffected by visual feedback (**Fig. 3E,F, S2H**), suggesting the presence of additional signals controlling this coordination. During straight walking, several animal species show compensatory mechanisms among eye, head and body that keep their gaze stable (Imai et al., 2001; Land, 1999). Notably, when flies walk forward, their slow body rotations are accompanied by anti-phase head movements to promote gaze stabilization (**Fig. S2**). Evidence for gaze stabilization has been previously reported in blowflies during walking (Blaj and Van Hateren, 2004), and in flight (van Hateren and Schilstra, 1999; Kern et al., 2006), although such anti-correlated head-body movement coordination had not been previously described. Several mechanosensory systems have been proposed to contribute to gaze stabilization (Bender and Dickinson, 2006b; Nalbach, 1993; Preuss and Hengstenberg, 1992). The presence of multiple signals suggests a functional analogy to gaze stabilizations reflexes found in vertebrates, such as vestibulo-ocular reflex (Waespe and Henn, 1987).

### Neural bases for straight course control

A simple, binocular visual model agent reproduces the exploratory behavior of real flies under ideal conditions (**Fig. 4**). However, signal dependent noise in motor and premotor systems (Harris and Wolpert, 1998), and speed-accuracy tradeoffs in motor control systems (Fitts, 1954; Meyer et al., 1988) typically render trajectories of movement variable. This adverse sensorimotor effect on the controller (Franklin and Wolpert, 2011) is further exacerbated by the external heterogeneities of the terrain. Under these higher noise conditions, multimodal, congruent interactions make the straightness performance more robust to noise, and comparable to real flies’ performance. We found concerted, congruent multimodal activity in HS and H2 cells (**Fig. 5**), two populations of cells that have long been proposed to be part of the flight course control system (Geiger and Nässel, 1981; Hausen, 1982b; Reichardt et al., 1983). This multimodal, congruent activity is consistent with the idea of a faithful representation of self-motion for robust course control, which may be based on the spanning of the dynamic range of the cells’ velocity tuning to high stimuli velocities (**Fig. 5**) (Chiappe et al., 2010), when compensatory movements may be more important (Fitts, 1954; Franklin and Wolpert, 2011). In addition, multimodal, congruent signals in HS and H2 cells may facilitate the proper interpretation of rotational flow signals by other postsynaptic networks (Bradley et al., 1996; Gu et al., 2008). Future experiments will focus on these important questions. Finally, our data also points to a functional organization of LPTC networks in the fly brain; HS and H2 cells, but not VS cells can drive leg-based angular movements of the fly (**Fig. 6**) (Busch et al., 2018; Fujiwara et al., 2017). This result is interesting because HS and H2 cells have separate projections from VS cells in a premotor area of the *Drosophila* fly brain (Namiki et al., 2018). Future work will examine whether there exists separate descending pathways connected to HS and H2 cells that control leg-vs. wing-based compensatory rotations (Schnell et al., 2017; Suver et al., 2016).

### A dynamic and selective modulation of neural activity by an internal signal can explain an asymmetry in the fly behavior

HS and H2 cells are modulated by a forward speed-related signal (**Fig. 5**). This signal is fast in both HS and H2 cells, suggesting a common origin related to, or highly correlated with, forward velocity. In nature, forward walking flies generate visual flow with translational and rotational components dominated by FTB visual motion, therefore driving activity in HS but not H2 cells. During forward segments, we found that the overall non-visual activity of H2 is decreased (**Fig. 5C**), and its direction selectivity diminished (**Fig. 5D**). Our data is consistent with the idea that during forward segments, H2 cells’ contribution to steering behavior is decreased, while HS cells’ contribution is enhanced (**Fig. 5,6**), revealing a matching between visual and non-visual activity that is dynamically modulated by an internal signal correlated to the forward velocity of the fly (**Fig. 7B**).

Under unilateral visual stimulation, when flies are exposed to either FTB or BTF rotational flow, we found that the reversed gain condition triggers different circling behavior (**Fig. 4**). In particular, our data indicates an asymmetry in the drift of angular velocity, which is prominent under FTB but small under BTF stimulation. One interesting speculation is that the sustained, reversed gain-induced angular velocity drift corresponds to the slow angular fluctuations under natural gain. Then, one possible explanation for the observed asymmetry in the angular velocity drift could result from the differential modulation FTB- and BTF-sensitive pathways by the forward velocity (**Fig. 5E**, **7B,C**). At high speed, when flies move forward, the FTB system triggers compensatory slow rotations, while the BTF system is inhibited. At low forward speed, both FTB and BTF systems could contribute to course control (**Fig. 7B**). We tested this idea by adding a forward velocity signal that differentially modulates the FTB and BTF elements of the visuomotor agent (see **Methods**). This modulation produces the asymmetry in the angular velocity drift observed under unilateral-stimulus reversed gain conditions in real flies (**Fig. 4B, C; 7C,D**). Previous studies have reported disparate results about the sensitivity of the fly to FTB vs. BTF direction of visual motion between walking and flight (Geiger, 1981; Götz and Wenking, 1973). We propose that these disparate results could be accounted for by the presence of an internal signal related to the forward velocity of the moving fly, differentially modulating the sensitivity of the fly to the different, self-generated directions of visual motion.

### Context-dependent visuomotor processing for locomotion control

Corollary discharge or efference copy is generally referred to an internal signal-based cancellation of sensory feedback (Crapse and Sommer, 2008; von Holst and Mittelstaedt, 1950; Straka et al., 2018). However, the brain actively uses reafference when computing the 3D structure of the world (Campbell et al., 2018; Kral, 2012; Schuster et al., 2002), or when required to monitor ongoing motor output (Chorev et al., 2016; Tschida and Mooney, 2012). Therefore, corollary discharge signals must be integrated within sensory feedback-related circuits in a context-dependent manner that is specified by the computations of the network. Our data shows that a network with course control function receives distinct velocity-related signals to tune the sensitivity of the circuit to visual feedback in a selective, dynamic and context-dependent manner. Importantly, the context is defined by the goal of the action, either to rapidly change course direction (Kim et al., 2017), or to stabilize it (**Fig. 5E, 7B**). The dynamic and context dependent control of the forward speed related signal suggests a predictive function on the network; the activity of neural elements that will not be excited by self-generated progressive visual flow is selectively tuned down, whereas the activity of those well situated to respond to such visual feedback is enhanced (**Fig. 5C-D, 7B**).

By taking advantage of the structure of exploratory behavior and analyzing the fly’s locomotion from a task-relevant perspective, we revealed a circuit mechanism for robust control of walking performance. Multisensory feedback dynamically interacts with internal, motor related signals to control behavior in a context dependent manner, a context that is set by motor commands or highly correlated internal signals. Such an internal control of sensorimotor circuits may be a fundamental mechanism to adaptively use sensory feedback information to guide locomotion in highly unpredictable natural environments. The numerical simplicity of the brain of *Drosophila melanogaster*, and the possibility to employ sophisticated genetic tools to target and manipulate the activity of identified cells, should allow establishing models for the functional relation between multimodal activity in recurrent networks and critical computations for motor control.

## Acknowledgements

We thank the Champalimaud Research’ fly facility for stock construction and maintenance, Kenta Asahina, Fernando Casares, Michael Dickinson, Gero Miesenböck, Michael Reiser, Gerald Rubin, and Marion Silies for kindly sharing flies. We are grateful to S. Huston, and Michael Reiser for early discussions about our results. We thank L. Petreanu, Marta Zlatic and members of the Chiappe’s lab for useful comments on the manuscript. This work was supported by the Champalimaud Foundation, by Foundação para a Ciência e a Tecnología FCT PD/BD/105947/2014 (T.C.), by the Japanese Society for the Promotion of Science JPSP 20170687 (T.F.), by the Bial Foundation grant 191/12 (M.E.Ch), by the Marie Curie Career Integration Grant PCIG13-GA-2013-618854 (M.E.Ch), and by the European Research Council Starting Grant ERC-2017-STG-759782 (M.E.Ch).

## Movie Legends

**Movie 1:** Left: Example exploratory path of a fly in a 5° Dot size visual environment. Right: Corresponding time series of the forward and angular velocities of the fly. Color code: saccades (blue) and forward segments (red) (see **Methods**).

**Movie 2**: Left: High resolution image of a freely exploratory walking fly labeled with 4 points used for head tracking using the DeepLabcut (Mathis et al., 2018). Right: Corresponding angular velocity (top) and head angle with respect to the body (bottom). Blue and red traces correspond to head angle based on DeepLabCut (blue), or based on a template matching strategy (red) (see **Methods**).

**Movie 3**: Exploratory paths in control flies, or in flies with the activity of T4/T5 cells silenced under natural or reversed gain conditions (see **Methods**).

## Methods

Further information and requests for resources and reagents should be directed to Eugenia Chiappe.

| REAGENT or RESOURCE | SOURCE | IDENTIFIER |
| --- | --- | --- |
| Antibodies |  |  |
| Mouse monoclonal anti-nc82 | DSHB | nc82, RRID:AB_2314866 |
| Chicken polyclonal anti-GFP | abcam | Cat#ab13970; RRID: AB_300798 |
| Rabbit polyclonal anti-GFP | Thermo Fisher Scientific | Cat#A6455; RRID: AB_221570 |
| Living colors DsRed polyclonal antibody | Clontech Laboratories, Inc. | Cat#632496; RRID: AB_10013483 |
| Alexa Fluor 488 conjugated goat anti-chicken | abcam | Cat#ab150169; RRID: AB_2636803 |
| Alexa Fluor 594 conjugated goat anti-mouse | Thermo Fisher Scientific | Cat#A11005; RRID: AB_2534073 |
| Alexa Fluor 633 conjugated goat anti-mouse | Thermo Fisher Scientific | Cat#A21050; RRID: AB_2535718 |
| Alexa Fluor 488 conjugated goat anti-rabbit | Thermo Fisher Scientific | Cat#A11008; RRID: AB_143165 |
| Alexa Fluor 594 conjugated goat anti-rabbit | abcam | Cat#ab150080; RRID: AB_2650602 |
| Alexa Fluor 594 Streptavidin | Thermo Fisher Scientific | Cat#S11227; RRID: AB_2313574 |
| Chemicals, Peptides, and Recombinant Proteins |  |  |
| Alexa Fluor 488 Hydrazide | Thermo Fisher Scientific | A10436 |
| Alexa Fluor 568 Hydrazide | Thermo Fisher Scientific | A10437 |
| Dextran, Texas Red™, 3000 MW, Lysine Fixable | Thermo Fisher Scientific | D3328 |
| Biocytin hydrazide | Thermo Fisher Scientific | B1603 |
| Collagenase IV | Worthington | CLS-4 |
| MgATP | Sigma | A9187 |
| Histamine | Sigma | H7125 |
| PFA | Electron Microscopy Sciences | 15710 |
| Normal goat serum | Life Technologies | PCN5000 |
| Experimental Models: Organisms/Strains |  |  |
| *Drosophila*: w<sup>1118</sup>; R59E08-p65ADZp;R42F06-ZpGdbd | Michael Reiser | N/A |
| *Drosophila*: w<sup>1118</sup>; Otd-nls:FLPo; + | Kenta Asahina | N/A |
| *Drosophila*: w<sup>1118</sup>; +; pJFRC56-10XUAS>myrtdTomato-SV40>eGFPKir2.1 | Gerald Rubin | N/A |
| *Drosophila*: UAS-2xEGFP | Bloomington Drosophila Stock Center (BDSC) | #6874 |
| *Drosophila*: w<sup>1118</sup>; 10XUAS-IVS-mCD8::RFP, 13XLexAop2-mCD8::GFP; +; + | BDSC | #32229 |
| *Drosophila*: w<sup>1118</sup>; +; VT058487-GAL4 | Vienna Drosophila Resource Center (VDRC) | N/A |
| *Drosophila*: w<sup>1118</sup>; +; 40XUAS-IVS-mCD8::GFP | BDSC | #32195 |
| *Drosophila*: w<sup>1118</sup>; +; UAS-P2X<sub>2</sub> | Gero Miesenböck | N/A |
| *Drosophila*: w<sup>+</sup>; UAS-Ort; + | Marion Silies | N/A |
| *Drosophila*: w<sup>1118</sup>; +; R81G07-GAL4 | BDSC | #40122 |
| *Drosophila*: w<sup>1118</sup>; R23C12-p65AD; + | BDSC | #70130 |
| *Drosophila*: w<sup>1118</sup>; +; R32A11-GAL4.DBD | BDSC | #69228 |
| *Drosophila*: w<sup>1118</sup>; 20xUAS-6xGFP/cyo; + | BDSC | #52261 |
| *Drosophila*: w<sup>1118</sup>; wg/cyo; 20xUAS-6xGFP | BDSC | #52262 |
| *Drosophila*: w<sup>1118</sup>; oddskipped-GAL4/cyo; + | Fernando Casares | N/A |
| *Drosophila*: w<sup>1118</sup>; +; R24E12-GAL4 | BDSC | #49084 |
| *Drosophila*: w<sup>1118</sup>; 8XLexAop2-FLPL; + | BDSC | #55820 |
| *Drosophila*: w<sup>1118</sup>; Cyo/Bl; UAS>stop>GtACR1/TM6B | this study | N/A |
| *Drosophila*: w<sup>1118</sup>; R39E01-lexA; + | BDSC | #52776 |
| *Drosophila*: 26XLexAop2-mCD8::GFP | BDSC | #32207 |
| *Drosophila*: w<sup>1118</sup>; R32A11-lexA/cyo; + | BDSC | #53564 |
| *Drosophila*: Top Banana wild type | Michael Dickinson | N/A |
| Software and Algorithms |  |  |
| MATLAB | The Mathworks, Natick, MA | N/A |
| ScanImage | Pologruto et al., 2003 | N/A |
| Ephus | Suter et al., 2010 | N/A |
| Visual Studio C# |  | Microsoft |

### Fly strains

*Drosophila melanogaster* flies were reared in standard medium at 25°C on a 12-hr light/12-hr dark cycle. Experiments were performed with 1 to 4 day-old male flies. We used the recently derived “Top Banana” WT line (TP) because they engaged in exploration more readily than other WT lines (**Fig.s 1**,**2**, **4** a gift from the Dickinson lab). To perturb neural activity, we used the ATP-gated cation channel P2X_2_ for activation. The inward rectified potassium channel Kir2.1, the light-gated anion channel GtACR1 (Mohammad et al., 2017), or the histamine-gated channel ort (Liu and Wilson, 2013) were used for inhibition. An intersectional approach designed with either Otd-nls:FLPo (Asahina et al., 2014), or 8XLexAop2-FLPL restricted the effector expression to the population of neurons of interest. The complete set of transgenic flies and the corresponding experiments are listed in Table S1. Specific sources of transgenic lines are listed in the Key Resources Table.

To generate 10X-*UAS-FRT-stop-FRT-GtACR1 (UAS>>GtACR1)* transgenic flies, the *Drosophila*-codon-optimized *GtACR1* sequence (Mohamamd et al, 2017) was subcloned into a pJFRC177-10x-*UAS-FRT-stop-FRT-*myr::GFP plasmid (addgene.org, plasmid #32149). The *GtACR1-EYFP* fragment from the p7-GtCAR1 plasmid was swapped in using CloneEZ® PCR Cloning Kit (GenScript) for the myr::GFP fragment in pJFRC177 and the sequence was verified (GenScript). The pJFRC177-10X-*UAS-FRT-stop-FRT-GtACR1* construct was injected into the *attP2* insertion site on the third chromosome (BestGene Inc.), and the transgenic progeny were balanced.

### Immunostaining

Isolated brains were fixed for 30 min at RT in 4% paraformaldehyde in PBS, rinsed in PBT (PBS, 0.5% Triton X-100 and 10 mg/ml BSA), and blocked in PBT + 10% NGS for 15 min. Brains were incubated in primary antibodies (1:25 mouse nc82 and 1:1000 rabbit antibody to GFP A-11120, Thermo Fisher Scientific) at 4 °C for three days. After several washes in PBT, brains were incubated with secondary antibodies (1:500 goat-anti rabbit: Alexa Fluor 488, A-11008, 1:500 goat-anti mouse: Alexa Fluor 633, A-21050 and 1:500 goat-anti mouse: Alexa Fluor 594, A-11032) for three days at 4 °C. Brains were mounted in Vectashield, and confocal images were acquired with a Zeiss LSM710 scope with a 40X oil-immersion objective.

### Virtual Reality setup for exploratory walking

Single flies with clipped wings moved freely in a 90mm circular arena with walls heated up to 42°C by an insulated nichrome wire (Pelican Wire P2128N60TFEWT). To prevent walking on the ceiling, the arena was covered with a glass plate pre-treated with Sigmacote (Sigma). The fly was video recorded using a near infrared camera (IDS UI-3240CP-NIR) at a frame rate of 60Hz and resolution of 900×900 pixels. The camera has attached a 2x extender (Computar EX2C), a wide field lens (Computar 5mm 1:1:4 ½), and a filter against visible light (Thorlabs AC254-100-A-ML). The illumination was set beneath the arena by IR LEDs (850nm SFH 4235-Z), powered by a current power supply (TENMA 72-10480). The position and orientation of the fly were determined *via* real time tracking. The tracking algorithm first performed a background subtraction using an image of an empty arena, and then applied a threshold. From the distribution of pixels within the contour of the threshold image, the 2D centroid and main orientation were estimated. The final estimate of the position and orientation of the fly was given by a Kalman filter, which combined the recorded data with prior knowledge about the system to minimize the difference between the real and estimated values statistically.

Visual stimuli were projected onto a rear projection material (Da-Lite High Contrast DA-Tex), attached to the floor of the arena. We used a small LED projector (DLP Lightcrafter 4500) at a frame rate of 60Hz and a pixel size of 160 px/mm. A custom-made software (FlyVRena, https://github.com/tlcruz/FlyVRena)(Cruz, 2013) generated the virtual worlds. FlyVrena tracks the position and orientation of the fly in real time, and uses computer game development libraries to render virtual objects, and update their behavior accordingly (closed loop). The delay between the fly movement and the update of world of ∼40-50ms (as measured using a photodiode, Vishay BPW21R) was largely generated by the projector system. Virtual worlds were 2D environments with sets of textured square-based random dots (**Fig.s 1-4, S1-S4**) or bars (**Fig. S3**). The models of the textured squares were designed in Blender (https://www.blender.org), the textures were generated in MATLAB and Adobe Illustrator, and rendered using the FlyVRena software.

Following FlyMad (Bath et al., 2014), high-resolution images were recorded at 120fps and 1024×544 pixel resolution with a near infrared camera (IDS UI-3360CP-NIR-GL) and a telecentric lens (Edmund Optics 1x, 220mm WD CompactTL Telecentric Lens). The camera pointed to a pair of galvo mirrors (Thorlabs GVS012/M - 2D Large Beam (10mm) Diameter Galvo System) that were moved based on the position of the fly to maintain the fly centered in the high-resolution frame. FlyVRena controlled the galvo mirrors.

### Head Tracking

High-resolution images were filtered with a Gaussian filter (width=16pixels), and background subtracted. Pixels that belonged to the fly were estimated with a combination of edge detector operations, and morphological transformations until a connected object with dimensions similar to the fly was detected. The centroid and orientation of the object were used to translate and rotate the frame in order to keep a vertical fly at the center of the frame. Next, a window around the head was cropped from the frame. The top of the thorax of the fly was masked out in the cropped frame, and the head was segmented using a combination of edge detector operations and morphological transformations. Head rotation was calculated via cross correlation with a template upright head (Berman et al., 2013). The head tracking error is the squared error between the template and the rotated head. The orientation that yielded the smallest error was kept as the real head orientation of the fly, with the associated angle defined as the head angle for that frame. The head angle trace was smoothed using a lowess algorithm with a window of 83 ms. Frames in which the head tracking error was high (top 2.5%) were removed from the original trace.

As an alternative strategy, we used DeepLabCut to estimate the rotation of the head with respect to the body of the fly (Mathis et al., 2018). We trained a neural network with a subset of images where the fly was in the upright position (see above) with four labels to track: Left Eye, Right Eye, Neck, and Thorax ending. After applying the tracking network to the full dataset, we transformed the positions of the labeled points into head angles. No significant differences were observed when comparing the two methods (Movie 2).

### Electrophysiological recordings

Details of the fly preparation for simultaneous physiology and behavior, and of the treadmill system are described elsewhere (Seelig et al., 2010). Briefly, a cold-anesthetized fly was mounted on a custom-made holder, and the back of the head’s cuticle was removed with fine tweezers. The dissected fly was mounted under the microscope and positioned on an air-suspended ball. *In vivo*, whole-cell patch-clamp recordings were performed using an upright microscope (Movable Objective Microscope, Sutter) with a 40× water-immersion objective lens (CFI Apo 40XW NIR, Nikon). The external solution, which perfused the preparation constantly, contained 103 mM NaCl, 3 mM KCl, 5 mM TES, 8mM trehalose, 10 mM glucose, 26 mM NaHCO_3_, 1 mM NaH2PO_4_, 4 mM MgCl_2_ and 1.5 mM CaCl_2_ (pH 7.3 when equilibrated with 95% O_2_/ 5% CO_2_; ∼275 mOsm). Patch pipettes (5–7 M?) were filled with internal solution containing 125 mM aspartic acid, 10 mM HEPES, 1 mM EGTA, 1 mM KCl, 4 mM MgATP, 0.5 mM Na_3_GTP, 20 µM Alexa 568–hydrazide-Na (Thermo Fisher Scientific, A10437) and 13 mM biocytin hydrazide (Thermo Fisher Scientific, B1603) (pH 7.3, final osmolarity: 260–265 mOsm).

The neural lamella was ruptured by local application of collagenase IV (Worthington)(Maimon et al., 2010). Current-clamp data was filtered at 4 kHz, digitized at 10 kHz using a MultiClamp700B amplifier (Molecular Devices), and acquired with Ephus (Suter, 2010). Unless otherwise stated, the recorded cell membrane potential (V_m_) was corrected for junction potential (11 mV).

To find H2-like cells, Texas Red Dextran (Thermo Fisher Scientific, D3328) was electroporated at the axon terminal of HS cells via brief voltage pulses (25ms pulses at 2Hz, amplitude: −10 to −50V, duration: 20-30s). While this strategy labeled different classes of neurons (data not shown), only one cell had contralateral dendritic projections in the second layer of the Lobula Plate (LP), a layer that houses regressive motion sensitive neurons (Maisak et al., 2013). In blowflies, the H2 and Hx cells possess such anatomy, but display a distinctive dendritic structure (Hausen, 1984; Krapp et al., 2001). Hx dendrites cover the entire surface of the LP, whereas H2 dendrites avoid the ventral and dorsal region of the LP. In *Drosophila*, a regressive-sensitive cell was previously identified as Hx in a transgenic line for the transcription factor *odd-skipped* (Levy and Larsen, 2013; Wasserman et al., 2015). We electroporated Texas Red at axon terminals of HS cells in a combined transgenic line (VT058487 and odd-Gal4) to compare the labeled cell with the odd-kipped expressing neuron **(Fig. S5Aii**). The cell soma clusters for HS/VS cells and for the odd skipped-expressing neurons were easily distinguishable (**Fig. S5Aii**, odd-skipped cells: gray arrowheads, and magenta arrow; HS/VS cells: green arrow). A single Texas-Red labeled cell with a large soma (∼ 10 μm) was co-labeled with GFP (**Fig. S5Aii**, magenta arrow). Examination of the dendritic structure of this neuron indicated an H2-like shape, with no ventral innervation of the LP. In addition, we identified a line from the Janelia collection (GMR32A11) with identical anatomy (**Fig. S5Aiii**). Both the Texas-Red labeled cell, and the LP neuron from GMR32A11 showed regressive visual motion selectivity (**Fig. S5B**). Because of the anatomical properties, and because physiology in the identified neuron gave always consistent results across all recorded cells, we referred to this cell as the *Drosophila* homologous H2 cell. Although the odd-skipped line may additionally labeled an Hx-like cell, the characterization of this transgenic line was out of the scope of this study. Recordings from HS and H2 cells were performed in darkness and under light conditions following Fujiwara et al., 2017.

### Generation of visual stimuli for electrophysiology

The visual display consisted of a 32×96 array of green LEDs (570 nm, Bet Lux Electronics (Reiser & Dickinson, 2008). The visual-field subtended angle was 180° in azimuth, and 50° in elevation (pixel size of 2.25°). Moving patterns were either sine-wave gratings with a constant spatial frequency (λ= 22.5°, **Fig. 5C, D, S5D**) or random dots of different dot size (2.25° to 9°, 16% occupancy, **Fig. S5B**). Visual response properties were characterized by moving the images horizontally at 22.5°/s for 3s along two opposing directions (3–5 repetitions for each condition). The stimuli in replay trials (**Fig. 5D**, **S5D**) were generated by the fly’s rotations in a closed-loop trial with a gain of 1.

### Unilateral activation of HS, H2, and VS cells

An electrode (5–7 M?) filled with ringer solution containing 10mM ATP (Sigma, A9187) and 50 µM Alexa Fluor 568 (Thermo Fisher Scientific, A10436) was placed in juxtaposition to the axon terminal of HS, H2 and VS cells, or dendrites of H2 cells. Pressure pulses (30 ms, 6 psi, picospritzer III, Parker) delivered ATP. For HS cells, we expanded the sample size to confirm previous observations (Fujiwara et al., 2017).

### Optogenetic silencing of HS cells

A fiber-coupled light (530 nm, M530F2, Thorlab) was projected through the objective lens onto one side of the brain, where the HS axons are located. Power density was calculated by the ratio between the power measured at the output of the objective, and the objective’s field of view. To dampen the illumination on the contralateral brain hemisphere, the head cuticle covering this side was kept intact. The neural activity manipulation was conditional to fly’s forward speed *via* a closed loop system. Real time treadmill signals were detected with a panel display controller unit (IO Rodeo) (Reiser and Dickinson, 2008). Once the forward velocity reached a threshold (> 1 mm/s on average for 3 s), a constant, 3s TTL pulse was sent to an LED driver (LEDD1B, Thorlab). We initially performed experiments at room temperature (24°C) and found that control flies not expressing GtACR showed a prominent startle response upon light stimulation. Increasing the bath temperature to 34°C largely decreased this response, suggesting that the light exerted a thermal effect. Higher bath temperature caused instability in whole-cell recordings; therefore, V_m_ changes upon light illumination were measured at room temperature.

### Chemogenetic silencing of HS cells

An electrode filled with external ringer solution with 1 mM histamine (Sigma) and 40 µM Alexa 568 was placed in juxtaposition to the axon terminal of HS cells. Brief pulses of histamine (10 ms, 6 psi) were applied conditioned to the fly’s forward speed (see above). Injection-triggered changes in V_m_ are shown only for those recordings that lasted until the end of the experiment (7/12 cells for experimental and 6/11 cells for control flies).

### Quantification and data analysis

We used MATLAB (MathWorks, Inc., Natick, MA) for data analysis.

### Data pre-processing for free walking assays

Post experiment, the position of the fly was transformed to forward and side velocities, whereas its orientation was transformed to angular velocity. Jump events of the fly were detected by a threshold in the fly acceleration (200mm/s2), and eliminated for subsequent analysis. Heading direction was inferred based on the persistence of the forward velocity, and if averaged forward speed between two jumps was negative, then orientation was rotated by 180° and velocities were re-calculated. All velocity signals on a window of 166ms around the jump were set to zero. The speeds were smoothed with a lowess algorithm with a window of 100ms. Activity was defined as translational speed greater than 0.5mm/s, or angular speed greater than 20°/s for at least 166ms or 10 video frames. On the other hand, inactivity bouts smaller than 333ms were considered activity. Walking bouts were a subgroup of activity bouts in which a sequence of movement lasted for at least 333ms (>3steps).

### Identification of spike-like events in the fly’s angular velocity

Spike-like events were fast and stereotyped rotations of the fly. To classify events as spike-like we used a continuous wavelet transform strategy (Arthur et al., 2013). For each walking bout, the continuous wavelet transform of the angular velocity was computed using Gaussian wavelets, and a signal with power at a range of 10-15Hz was extracted (frequency signal). Note that the selection of the frequency band affects the spike detection, but it is not a determinant factor in subsequent analysis (**Fig. S1**). Next, the local maxima of the frequency signal, and of the absolute value of the angular velocity were calculated. If these local maxima coincided between the two sets of local maxima, these were labeled as putative spike turns (**Fig. S1A**).

Putative spike turns were winnowed twice to remove small local maxima (<200°/s) that did not disrupt the forward velocity of the fly (variance of forward speed < 3mm/s), and to match the signal with a template obtained via PCA on a subset of pronounced spike-like events (the first principal component explained ∼90% of the shape variance, and was used as the template). Putative spike turns in which the square distance between the scaled shape and the template was smaller than a cutoff of 0.15 were labeled as spike-like events, which were simply referred to as body saccades throughout the main text (**Fig. S1A**). The free parameters of the classifier did not affect the results over a wide range of possible parameters (**Fig. S1B**). We also observed that body saccades were heavily modulated by the presence of the hot walls, both in magnitude (**Fig. S2D**) and direction (data not shown).

To identify forward walking segments with no body saccades, spike-like events were removed from the walking bout. The remaining segments longer than 333ms, and with average forward velocity larger than 6mm/s were defined as forward segments throughout this study.

### Straightness and visual influence

A 333ms window centered on each point of a forward segment path was used to calculate the distance between the data points and a line defined by the edges of the window (deviation from an ideal straight path). Straightness per forward segment was defined as the sum of traveled distances within the window divided by the sum of deviations. Straightness per fly is the weighted average of all the straightness per forward segment, with weights given by the total distance walked in each segment.

Under natural gain conditions, when a fly rotates to the left, the world rotates to the right. In contrast, under the reversed gain condition, when the fly rotates to the left, the world rotates left too. The reversed gain condition induces a persistent rotation of the fly, a behavior known as circling (von Holst and Mittelstaedt, 1950). Visual influence was defined as the difference between the probabilities of circling in reversed vs. natural gain conditions (**Fig. S3**). In darkness or for 1° size dots, the probability to rotate in the same or opposite direction under natural gain conditions is very similar, and therefore, due to the intrinsic variability of the measurement, in some cases it was possible to obtain a negative visual influence scalar. Under the dark condition we artificially divided the dataset into the same trial structure as in the random dot stimulus experiment and calculated visual influence in the same way. Visual influence was calculated either for the full time series, or during forward segments only.

Due to the delay between the estimate of the fly’s position and orientation and the update of the visual environment, when the fly stopped walking under the reversed gain condition, the visual environment continued rotating with the previous angular velocity of the fly for three consecutive frames. This brief “open-loop” segment could induce a head optomotor response. Typically, after the stimulus stop moving, the head remained with an offset position driven by the stimulation until the fly initiated a behavior

### Simulations

A simulated rigid-body agent moved forward or executed a saccade in alternating activity and immobility intervals (**Fig. S4A, B**). The distribution of the length of activity bouts was similar to the real flies’ distribution. The agent explored a simulated 90mm circular arena with hot walls. Saccades were modeled as highly stereotyped rapid events in the angular velocity of the agent (**Fig. S2B**). The probability and amplitude of a saccade depended on the distance to the arena wall. The time intervals within an activity bout without saccades were considered forward segments. Forward segments had constant translational speed (20 mm/s), and non-zero slow rotations represented by an “injected” 1/f noise. The size of the noise was scaled by the parameter ‘Noise Level’ (**Fig. S4C**). In the full version of the model, there were two rotational control loops (**Fig. 4Eii**). The first one was based on a copy of the angular velocity of the agent of the previous 50ms (motor feedback), which was filtered by a temporal kernel (Fujiwara et al., 2017) and scaled by the parameter ‘Motor Weight’ to subtract it from the current angular velocity. The second rotational controller was based on visual flow. Four channels containing arrays of a “two-quadrant” type Hassenstein-Reichardt (HR) correlator detected self-generated visual motion cues (Eichner et al., 2011). Self-generated visual stimuli were modeled first by high-pass filtering the visual pattern at each spatial position (τ = 50ms). This was followed by a half-wave rectification step. Next, the visual signal was low-pass filtered (first order, τ = 15ms), and then multiplied with an unfiltered signal from a neighboring spatial location. This was done twice in a mirror-symmetrical manner, followed by subtraction, yielding a fully opponent direction-selective output signal, which was then scaled by the parameter ‘Visual Weight’ to subtract it from the future rotational speed (compensatory rotation).

To find the model that most approximated the behavioral data, we started by simulating data in the dark using various noise levels and motor weights. We selected the set of free parameters (noise, motor weight and visual weight) such that the agent approximated the behavior of a fly. In these models the relation between motor weight and noise level was linear (**Fig. S4C** inset), thereby reducing the amount of free parameters for the second set of simulations. We then simulated the agent with various levels of visual weight and noise level under different visual environments. For each model, the distance between the simulated and the measured straightness was calculated for all visual environments. The model with the smallest distance to the data was selected as the best fitted model.

Finally, for the simulations in (**Fig. 7**), each visuomotor channel was modulated by the forward speed of the agent. Specifically, the BTF channels output were divided by whereas the FTB were multiplied by a fifth of the forward speed value. The unilateral stimulation was identical to the real fly’s unilateral stimulus configuration.

### Data analysis for electrophysiology

Walking-related signals in HS and H2 cells were analyzed following (Fujiwara et al., 2017). Briefly, we extracted walking bouts from the treadmill signals using a supervised machine-learning algorithm JAABA (Kabra et al., 2013) based on side-view videos of the walking fly (100 Hz, A602f, Basler). Isolated walking bouts, and the corresponding V_m_ signals (for spiking neurons, sub-threshold signals were inferred by clipping spikes within a ±5 ms window, and interpolating the original traces) were concatenated per fly. All signals were down-sampled to 500 Hz and smoothed using a lowess algorithm with a 120ms window. For **Fig. 5B**, V_m_ was triggered when the angular velocity exceeded an absolute value of 200 °/s. For **Fig. S5D**, the V_m_ was projected onto a 2D behavioral map (angular velocity bins: 20 °/s vs. forward velocity bins: 0.5 mm/s), or 2D visuomotor map (visual velocity bins: 20 °/s vs. angular velocity bins: 20 °/s). Each pixel of the map was defined as the mean of all collected V_m_ values at the corresponding parameter space combination.

To infer direction selective responses (DS responses), V_m_ was normalized to the maximum value per cell. For congruent combinations of visual and non-visual signals (**Fig. 5C, D**), DS were defined as the absolute value of the difference between the mean pixel values for the preferred vs. null visual direction selectivities within the congruent quadrants of the visuomotor map (**Fig. S5D**). DS responses for opposite combinations of visual and non-visual signals were calculated as the absolute value of the difference between the preferred vs. null visual direction selectivities within the opposite quadrants of the visuomotor map (**Fig. S5D**). Vision-only DS responses were calculated at moments when flies were stationary. Rotation-only DS responses were calculated as the absolute value of the difference between the preferred vs. null motor direction selectivity when the visual display was stationary.

### Statistics

We performed two-sided Wilcoxon signed-rank test for paired groups, two-sided Wilcoxon rank-sum (here, abbreviated as MWW) test for comparisons between two independent groups, t-test for the correlation analysis, and two-way ANOVA followed by Tukey-Kramer test for multiple comparisons. In addition, for statistics on slopes in **Fig. 5F**, we performed a resampling, bootstrapping based method, were 1% of all data was randomly chosen to fit a linear regression of slopes and offsets for low vs. high forward velocity. This procedure was repeated 1000 times to obtain the distributions for the difference in slope or offsets for fits obtained in low vs. high speed, with zero representing no difference. P-values represent the probability that the slopes (or offsets) of the linear regressions for bootstrapped data obtained in “low” vs. “high” speed are equal.

